# Nicked tRNAs are stable reservoirs of tRNA halves in cells and biofluids

**DOI:** 10.1101/2022.08.31.506125

**Authors:** Bruno Costa, Marco Li Calzi, Mauricio Castellano, Valentina Blanco, Ernesto Cuevasanta, Irene Litvan, Pavel Ivanov, Kenneth Witwer, Alfonso Cayota, Juan Pablo Tosar

## Abstract

Nonvesicular extracellular RNAs (nv-exRNAs) constitute the majority of the extracellular RNAome, but little is known about their stability, function and potential use as disease biomarkers. Herein, we measured the stability of several naked RNAs when incubated in human serum, urine and cerebrospinal fluid (CSF). We identified extracellularly produced tRNA-derived small RNAs (tDRs) with half-lives of up to three hours in CSF. Contrary to widespread assumptions, these intrinsically stable small RNAs are full-length tRNAs containing broken phosphodiester bonds (i.e., nicked tRNAs). Standard molecular biology protocols, including phenol-based RNA extraction and heat, induce the artifactual denaturation of nicked tRNAs and the consequent in vitro production of tDRs. Broken bonds are roadblocks for reverse transcriptases, preventing amplification and/or sequencing of nicked tRNAs in their native state. To solve this, we performed enzymatic repair of nicked tRNAs purified under native conditions, harnessing the intrinsic activity of phage and bacterial tRNA repair systems. Enzymatic repair regenerated an RNase R-resistant tRNA-sized band in northern blot and enabled RT-PCR amplification of full-length tRNAs. We also separated nicked tRNAs from tDRs by chromatographic methods under native conditions, identifying nicked tRNAs inside stressed cells and in vesicle-depleted human biofluids. Dissociation of nicked tRNAs produces single-stranded tDRs that can be spontaneously taken up by human epithelial cells, positioning stable nv-exRNAs as potentially relevant players in intercellular communication pathways.

## INTRODUCTION

Extracellular RNAs (exRNAs) circulating in human bodily fluids can enable disease diagnosis before the onset of clinical symptoms (Moufarrej et al., 2022). Beyond applications as disease biomarkers in liquid biopsies (Heitzer et al., 2019), exRNAs are also involved in intercellular communication pathways between cells in different tissues (Thomou et al., 2017) and in host-pathogen interactions (Buck et al., 2014; Cai et al., 2018; Garcia-Silva et al., 2014).

One key aspect governing both exRNA functionality and utility as biomarkers is their stability against ubiquitous extracellular RNases (Tosar et al., 2021). This can be achieved by RNA encapsulation inside extracellular vesicles (EVs) such as exosomes and microvesicles (Skog et al., 2008; Valadi et al., 2007). However, despite the functional relevance (Mateescu et al., 2017) and biotechnological applications (O’Brien et al., 2020) of EV-encapsulated RNA, the majority of exRNAs in cell culture (Tosar et al., 2015; Turchinovich et al., 2011; Wei et al., 2017; Zhang et al., 2021) and in human plasma (Arroyo et al., 2011; Geekiyanage et al., 2020; Turchinovich et al., 2011; Vickers et al., 2011) are not transported as part of EV cargo.

Transfer RNA-derived RNAs (tDRs) are among the most abundant nonvesicular small RNAs (nv-exRNAs) in cell culture (Tosar et al., 2015; Wei et al., 2017). Inside cells, tRNA cleavage and the consequent generation of specific tDRs is a conserved response to stress in all kingdoms of life (David et al., 1982; Li et al., 2008; Thompson et al., 2008). In humans, RNase A superfamily members, such as RNase 5 (Angiogenin), are responsible for stress-induced tRNA cleavage at the anticodon, generating stress-induced tRNA halves (tiRNAs) (Fu et al., 2009; Yamasaki et al., 2009). tiRNAs can regulate gene expression at various levels, including global inhibition of translation initiation (Ivanov et al., 2011; Lyons et al., 2020). Other, shorter tDRs can bind to mRNAs and regulate their translation (Kim et al., 2017) or silence genes by a miRNA-like mechanism (Kuscu et al., 2018).

Extracellular tDRs were first reported in EVs from murine immune cells (Nolte’T Hoen et al., 2012) but were later shown to be present mainly outside vesicles in mouse serum (Dhahbi et al., 2013; Zhang et al., 2014). In human breast cancer cell lines, tDRs can be detected in vesicular fractions but are overwhelmingly more abundant in EV-depleted ultracentrifugation supernatants (Tosar et al., 2015). While a heterogeneous population of tDRs is detectable inside cells, extracellular nonvesicular tDRs are mainly 5’ tRNA halves of 30 or 31 nucleotides, derived from tRNA^Gly^ and tRNA^Glu^. These specific fragments are also ubiquitous in human biofluids (Srinivasan et al., 2019). A possible explanation for the extracellular enrichment of these fragments is their enhanced stability against degradation, because 5’ tRNA^Gly^_GCC_ halves of 30 – 31 nt can form RNase-resistant homodimers in vitro (Tosar et al., 2018).

To study exRNA processing in more detail, we added RNase inhibitors (RI) to cell-conditioned media and uncovered a population composed of nonvesicular ribosomes and full-length tRNAs (Tosar et al., 2020). Knock-out of RNase 1 in K562 cells also shaped exRNA profiles from tRNA halves into full-length tRNAs (Nechooshtan et al., 2020). Thus, most nonvesicular tRNA halves are generated directly in the extracellular space by endonucleolytic cleavage of extracellular full-length tRNAs (Nechooshtan et al., 2020; Sanadgol et al., 2022; Tosar et al., 2020).

We have found that extracellular ribosomes can induce dendritic cell activation in vitro in an exRNA-dependent manner (Tosar et al., 2020). However, the involvement of nonvesicular exRNAs (nv-exRNAs) in intercellular communication pathways faces an important conceptual challenge. As a consequence of the strong ribonuclease activity that characterizes the extracellular space (Sorrentino, 2010), nv-exRNAs are expected to be rapidly degraded unless protected by RNA-binding proteins. How, then, do these RNAs resist degradation, diffuse to recipient cells and trigger downstream effects, or even remain measurable and thus serve as potential disease biomarkers?

In this study, we incubated protein-free RNA or purified ribonucleoprotein complexes in human biofluids and measured their decay kinetics by northern blot. As expected, the protective effect of RNA-binding proteins was confirmed. Interestingly, though, we also found that certain specific naked tRNAs are intrinsically stable. Furthermore, the stability of 5’ tRNA halves generated from the endonucleolytic cleavage of unstable tRNAs was extremely high, with half-lives of hours in certain human biofluids. Also surprisingly, these ultra-stable RNAs are not single-stranded. Developing and employing different strategies of enzymatic repair and electrophoretic or chromatographic separations under native conditions, we show that these RNAs natively exist as full-length tRNAs containing broken phosphodiester bonds. Commonly used RNA purification methods, sequencing, and denaturing northern blotting do not detect these forms, forcing them to melt into single-stranded tRNA halves. Nevertheless, at low rates, nicked tRNAs also seem to disassemble spontaneously into tRNA halves in the extracellular space. These extracellular nonvesicular tDRs are unstable, but they can be efficiently internalized by recipient cells.

## RESULTS

### Identification of naked RNAs that are stable in the presence of serum

To search for intrinsically stable RNAs in extracellular samples, we first screened the most abundant cellular transcripts to see if, in the absence of their protein counterparts, they can resist degradation in serum-containing media (**Figure 1A)**. Unsurprisingly, naked rRNAs were degraded in less than one minute as judged by lack of northern blot signals in the absence of added RNase inhibitors (RI). In contrast, and consistent with our previous observations (Tosar et al., 2020), full-length tRNA^Lys^_UUU_ was present at input levels after 1.5 hours, suggesting that this tRNA is not efficiently targeted by serum RNases. This was not observed for other tRNAs like tRNA^Gly^_GCC_, tRNA^Asp^_GUC_ and tRNA^Glu^_GUC_ which, like rRNAs, were undetectable after one minute (**Figure 1A** and data not shown).

**Figure 1:**
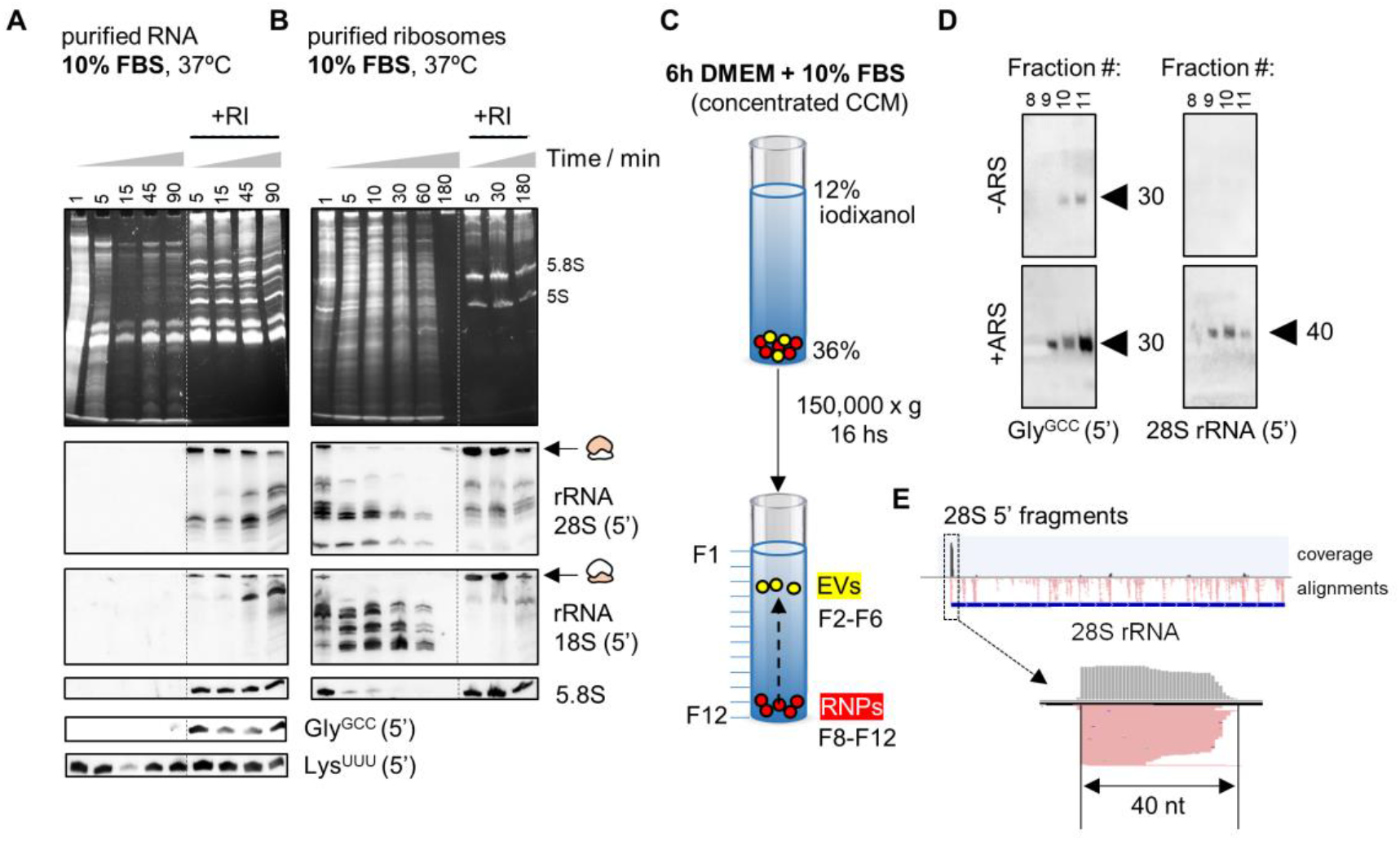
Identification of stable nonvesicular RNAs. Northern blot of different rRNAs and tRNAs after incubating purified RNA (A) or ribosomes (B) from human cells in 10% FBS. C-D) Northern blot of 5’ tRNA^Gly^_GCC_ (D, left) or 5’ 28S rRNA-derived fragments (D, right) in extracellular nonvesicular fractions purified by density gradients (C). U2-OS cells were treated or not with NaAsO_2_ (ARS) before collecting the cell-conditioned medium (CCM). E) Read coverage in small RNA-seq data of extracellular ribosomes (from Tosar et al., 2020), revealing enrichment of 40 nt 5’-derived small RNAs among all other 28S rRNA-derived fragments.

### Ribosomal proteins stabilize rRNAs and protect rRNA-derived fragments

To assess the effect of RNA-binding proteins on the stability of nv-exRNAs, we purified whole ribosomes from cells (**Figure S1**) and studied rRNA decay kinetics in 10% FBS (**Figure S1B**). In ribosomes, full-length rRNAs and rRNA-derived fragments were detectable for at least 10 and 60 minutes at 37ºC, respectively, even in the absence of RI. Thus, ribosomes are more stable than naked rRNAs and ribosomal proteins greatly stabilize extracellular rRNA-derived fragments.

Based on these results, we searched for nonvesicular rRNA-derived fragments in cell-conditioned medium (CCM) obtained in the presence of 10% FBS. We separated EVs from RNP complexes by flotation on iodixanol density gradients (**Figure 1C**). Extensive characterization of this methodology has shown that the densest fractions (8-12) are devoid of EVs (Jeppesen et al., 2019; Tosar et al., 2020). We reproduced our previous findings (Tosar et al., 2020) showing the presence of 5’ tRNA^Gly^_GCC_ halves of 30 – 31 nt and the absence of full-length tRNAs in these nonvesicular fractions (**Figure 1D**). Interestingly, northern blot detection of these fragments was enhanced when cells were treated with the cytotoxic agent sodium arsenite (**Figure 1D**, +ARS). Additionally, using a probe complementary to the first 20 nucleotides of the 28S rRNA resulted in a single band of 40 nt in the conditioned medium of arsenite-treated cells. Purified extracellular ribosomes analyzed by small RNA-seq (Tosar et al., 2020) also showed enrichment of these 40 nt 5’ fragments (**Figure 1E**). Detection of these fragments in the presence of serum suggests that at least some stable RNA-containing RNPs are produced after the release and subsequent fragmentation of extracellular ribosomes, in agreement with recent findings (Tosar et al., 2022).

### Naked full-length tRNA^Lys^_UUU_ is intrinsically stable in biofluids

To precisely measure tRNA half-lives, we repeated the previous assay with better temporal resolution (**Figure 2A**). Strikingly, the half-life of tRNA^Lys^_UUU_ (∼750 s) was 58-fold greater than that of tRNA^Gly^_GCC_ (∼13 s) and > 125-fold greater than that of rRNAs or the 7SL RNA (< 6 s). In vitro incubation of RNA with recombinant human RNase 1 (r-RNase1), representing the most common RNase in human blood, resulted in virtually identical results (**Figure 2A**). Thus, naked tRNA^Lys^_UUU_ is intrinsically resistant to the action of RNase A family members.

**Figure 2:**
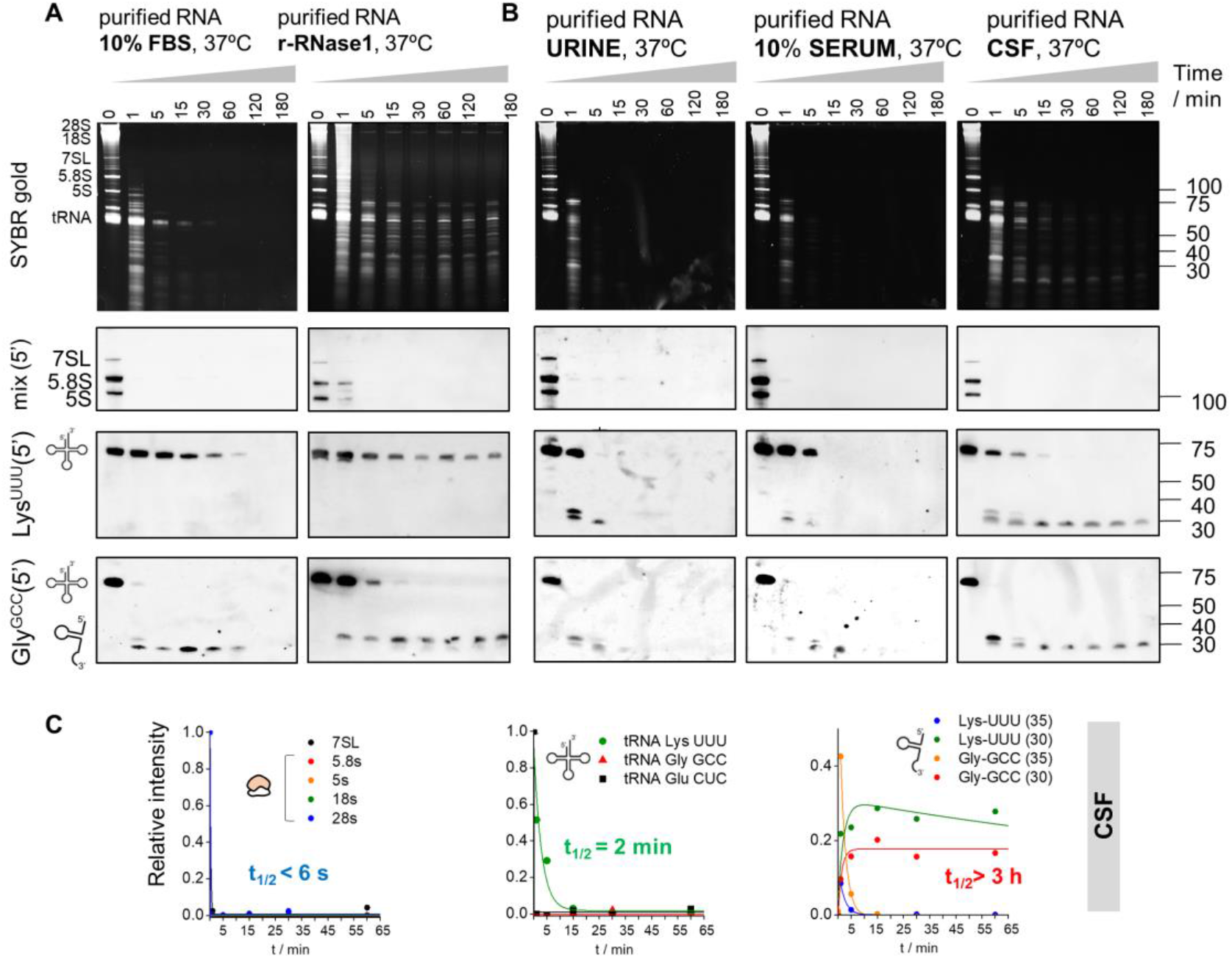
Naked tRNA halves are extremely stable in human biofluids. Northern blot of several noncoding transcripts after incubating purified RNA from human cells in 10% FBS or with recombinant human RNase 1 for different periods (A). Samples were also incubated in human urine, 10% serum and CSF (B). Half-lives in CSF were calculated for all tested RNAs and shown in (C). r-RNase1: recombinant human RNase 1.

We then repeated these assays in human biofluids including urine, diluted and undiluted serum, and cerebrospinal fluid (CSF) (**Figure 2B** and **Figure S2A**). Strikingly, the stability of full-length tRNA^Lys^_UUU_ was always higher than the stability of any other tested RNA, irrespective of the sample type (**Figure 2B, Figure S2B** and **S2C**).

### Glycine tRNA halves produced in biofluids are extremely stable

Full-length tRNA^Gly^_GCC_ was almost completely degraded in less than one minute in 10% FBS (**Figure 2A**) and in human biofluids (**Figure 2B**). However, its cleavage resulted in the formation of 5’ halves that showed remarkable long half-lives, even in undiluted human sera (**Figure 2A, 2B** and **Figure S2**). For instance, the half-life of 5’ tRNA^Gly^_GCC_ halves of 30 nt in CSF was longer than 3 hours (**Figure 2C**), meaning that these RNAs are highly stable once they are generated and will therefore persist in biofluids.

A closer inspection of the data shows strong differences in the stability of fragments derived from the same parental tRNAs but with slightly different lengths. Consistent with our previous report (Tosar et al., 2020), tRNA^Gly^_GCC_ is first cleaved at the anticodon loop, generating 34 – 35 nt 5’ halves that rapidly disappear. These fragments are subsequently replaced by highly stable shorter fragments of approximately 30 – 31 nt (t_1/2_ = 327, 439 and > 10,800 s in urine, diluted serum and CSF, respectively, **Figure 2B**), with a cleavage site at the start of the anticodon loop. An additional cleavage site in the TΨC loop (position 54) could be identified and exposed by using lower r-RNase1 concentrations (**Figure S3A**).

The sequential production of first longer (34 – 35 nt) and then shorter (30 – 31) 5’ halves from tRNA^Gly^_GCC_ was also observed with recombinant human RNase 5 or Angiogenin (r-Ang; **Figure S4**). Thus, cleavage sites and tRNA processing dynamics seem to be conserved among different RNase A family members.

Overall, tRNAs are more resistant to degradation than other longer noncoding RNAs, but differences in stability among tRNAs (e.g., tRNA^Lys^_UUU_ vs tRNA^Gly^_GCC_) are also substantial (> 50-fold). Consistent with our previous reports (Tosar et al., 2018, 2020), naked 5’ tRNA^Gly^_GCC_ halves of 30 – 31 nt are accumulated at higher RNase concentrations or after longer incubations in biofluids and are extremely stable.

### Nicked tRNAs are a source of 5’ and 3’ tRNA halves

Next, we compared the relative stabilities of 5’ and 3’ tRNA^Gly^_GCC_-derived fragments (**Figure S5**). Surprisingly, 30 – 35 nt 3’ fragments were observed, and their rate of decay was comparable to their 5’ counterparts. In certain biofluids, such as FBS (**Figure S5A**) and urine (**Figure S5B**), 3’ tRNA-derived fragments < 20 nt were observed after short incubations (1 min; black asterisks). However, these fragments presented a very short half-life, in sharp contrast with the 5’ and 3’ halves.

We considered the possibility that 5’ and 3’ tRNA^Gly^_GCC_ halves could remain physically associated with each other after RNase cleavage, representing a full-length tRNA bearing a cleaved phosphodiester bond at the anticodon loop (i.e., in the form of “nicked tRNAs”). This would explain the similar decay kinetics of each half among different biofluids. The introduction of any irreversible denaturation steps in analytical determinations, such as those used in standard molecular biology approaches, would induce dissociation of nicked tRNAs into single-stranded tRNA halves. Therefore, testing this hypothesis required the generation of new assays capable of interrogating oligomeric RNA complexes under native conditions.

Nicked tRNAs are the natural substrates of T4 polynucleotide kinase (PNK) and T4 RNA ligase 1 (Rnl1) (Schwer et al., 2004). When *Escherichia coli* is infected by the T4 Phage, a bacterial anticodon nuclease is activated and cleaves the host’s tRNA^Lys^ in an attempt to prevent the translation of viral proteins (Kaufmann, 2000). This results in a nicked tRNA bearing a 3’ cyclic phosphate (3’ cP) and a 5’-OH adjacent to the cleavage site. However, the phage evolved two enzymes capable of performing end-healing (PNK) and tRNA repair (Rnl1). We harnessed the intrinsic activity of these enzymes and used them to investigate the native structure of human tRNA halves in extracellular samples.

To optimize the system, we purified total RNA from cells and incubated the RNA with r-RNase1 for 0, 15 or 60 minutes (**Figure 3A**). As expected, by 60 minutes, full-length tRNA^Gly^_GCC_ was completely degraded and converted to a collection of tDRs (**Figure 3A**). RNase 1 degradation products were purified by phenol-free silica-based solid phase extraction (SPE) columns and treated with either PNK alone, PNK followed by Rnl1, or PNK followed by Rnl2 (a dsRNA-specific ligase). Surprisingly, treatment with PNK and Rnl1 regenerated a single band of approximately the size of the cognate full-length tRNA (**Figure 3A** and **3B**). Ligation with Rnl2 was less efficient.

**Figure 3:**
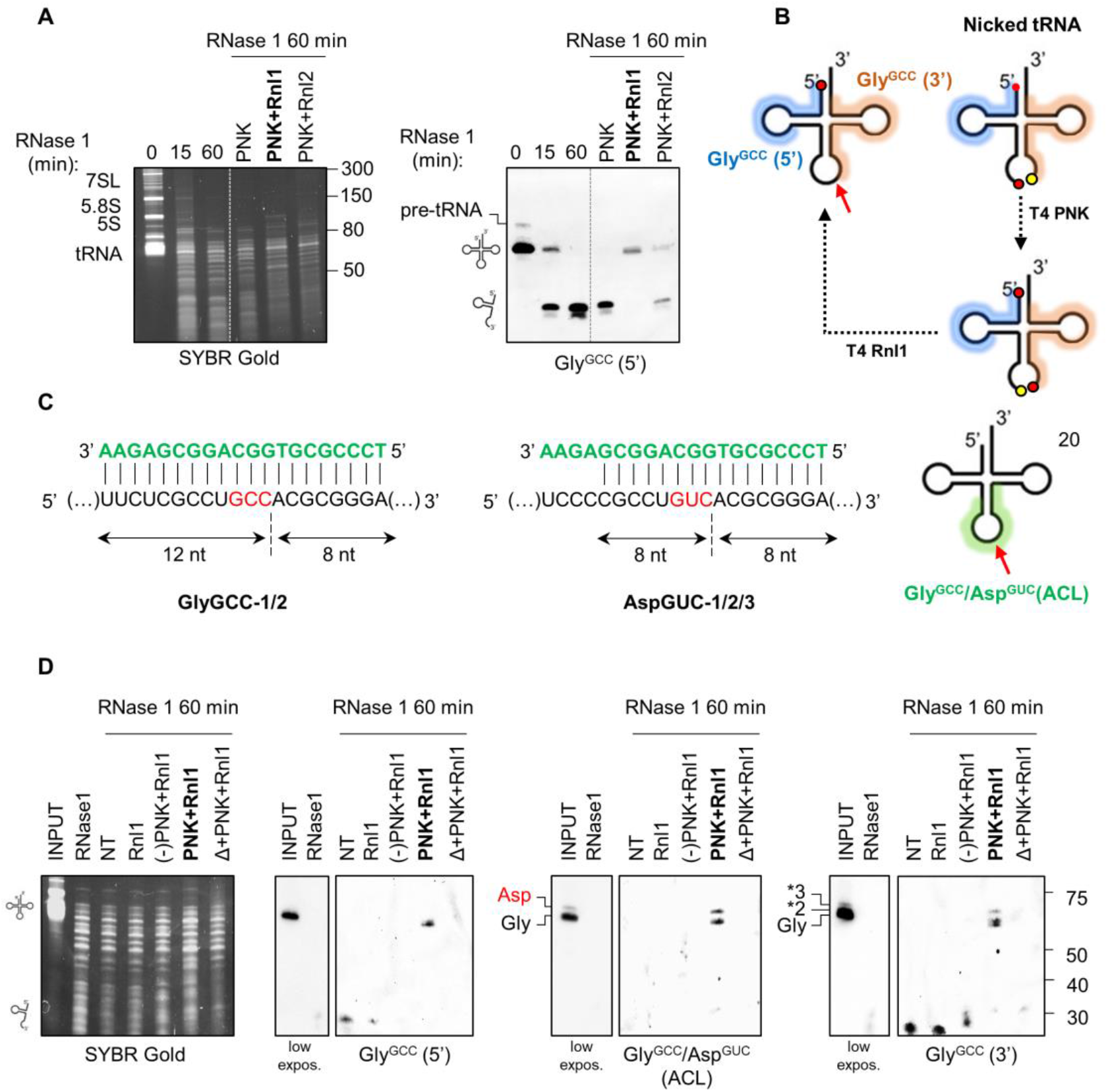
Most tRNA halves identified by northern blot are nicked tRNAs. A) RNase 1-treated RNA (60 min) was incubated with the indicated enzymatic combinations. Repair of tRNA^Gly^_GCC_ was analyzed by northern blot using a 5’-targeting probe. B) schematic representation of the nicked tRNA repair strategy and 5’ (blue) and 3’ (orange) probe binding sites in tRNA^Gly^_GCC_. C) Design of a third probe targeting the anticodon loop (ACL) of tRNA^Gly^_GCC_ and tRNA^Asp^GUC. D) enzymatic repair of tRNA^Gly^_GCC_ evidenced with either the 5’, ACL or 3’ probes. NT: fragmented RNA purified by SPE but without treatment with the enzymatic repair cocktail. (-)PNK: mutant PNK lacking phosphatase activity. Δ: heat.

To demonstrate that the tRNA-sized re-ligated products are indeed repaired tRNAs, we designed a third probe (termed ACL, for anticodon loop) bridging both sides of the anticodon of tRNA^Gly^_GCC_ (**Figure 3C**). This probe was designed so that the T_m_ of its pairing with either the 5’ or the 3’ half was below the hybridization temperature of the assay (42ºC). Thus, the ACL probe would fail to detect 5’ or 3’ tDRs but should be able to hybridize with full-length or repaired tRNA^Gly^_GCC_. Due to sequence similarities among tRNAs, ACL also recognizes the anticodon loop of tRNA^Asp^GUC, but fortunately, these tRNAs migrate slightly differently in denaturing urea gels, making this assay multiplex. Strikingly, treatment of RNase 1-treated RNA with PNK and Rnl1 regenerated tRNA-sized products observable with the 5’, the 3’ or the ACL probes (**Figure 3D**). Furthermore, heating (Δ) and then cooling the RNase-treated RNA before the addition of the enzymatic cocktail (Δ+PNK+Rnl1) prevented the generation of a tRNA-sized band, demonstrating that ligation occurres in *cis* under the assay conditions. Rnl1 alone or following incubation with a mutant version of T4 PNK lacking its 3’ phosphatase activity [(-)PNK] also failed to reconstitute a full-length tRNA.

Next, we studied whether tRNA halves generated in human biofluids (**Figure 2B**) are in fact nicked tRNAs. Indeed, a short incubation in CSF (1 min) produced 5’ tRNA^Asp^GUC halves that could be enzymatically repaired back into a full-length tRNA (**Figure S6A**). Enzymatic repair of tRNA^Gly^_GCC_ produced after longer (1 hour) incubations in CSF was also highly efficient (**Figure S6B**). However, the main re-ligated product was approximately 11 nt shorter than the parental full-length tRNA, consistent with an irreversible loss of the entire anticodon loop (7 nt) plus the single-stranded NCCA 3’ overhang.

In summary, nicked tRNAs are produced in vitro and in biofluids using different tRNAs as substrates. These nicked tRNAs can be repaired enzymatically to regenerate an almost full-length tRNA, presumably lacking the 3’ NCCA overhang in in vitro settings (Akiyama et al., 2022; Nechooshtan et al., 2020). Additional bases can be trimmed after prolonged exposure to extracellular RNases, but these shorter forms are still reparable using our assay.

### Nicked tRNAs are irreversibly melted by popular RNA extraction methods

Whereas heating RNase-treated RNA before tandem incubation with PNK and Rnl1 prevented the formation of a tRNA-sized band, this also affected the detection of monomeric 5’ and 3’ tRNA halves (**Figure 3D**; Δ+PNK+Rnl1 lane). However, when we repeated SPE purification under conditions optimized for small RNA recovery (**Figure S6C**), monomeric tRNA halves were now detectable in the control (i.e., heat) reaction (**Figure 4A**; red arrows).

**Figure 4:**
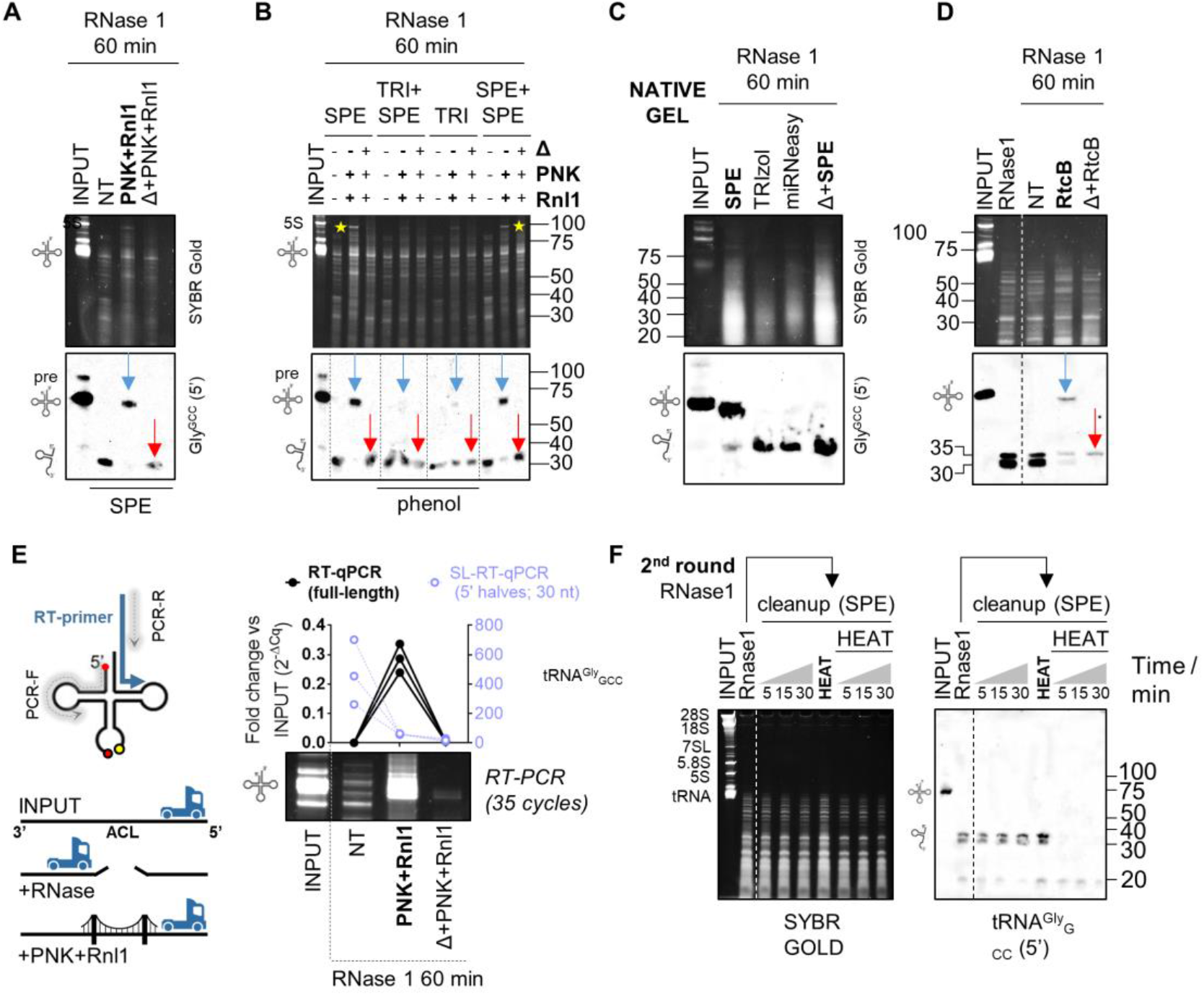
Nicked tRNAs are stable, dissociated by phenol, can be repaired by PNK+Rnl1 or by RtcB, and cannot be reverse-transcribed otherwise. A) Enzymatic nicked tRNA repair under optimized SPE conditions. Blue arrow: repaired product (ligation in cis). Red arrow: single-stranded tRNA halves generated by heat before enzymatic treatment. B) RNA purification by TRIzol (TRI) impairs enzymatic nicked tRNA repair. Yellow stars: other RNAs are also repaired and ligated in cis. C) Northern blot of RNAse1-treated RNA purified by SPE, TRIzol, miRNeasy or heated before SPE purification, after separation in native gels. D) Nicked tRNA repair with RtCB from *E. coli*. E) RNase treatment generates a roadblock for thermostable reverse transcriptases (RT) at the anticodon loop (ACL) that inhibits RT-PCR of nicked tRNAs. Treated samples were analyzed by end-point RT-PCR (SYBR gold-stained gel) or RT-qPCR for the full-length tRNA^Gly^ GCC (black, left axis) or by SL-RT-qPCR for 5’ tRNA^Gly^ GCC halves (purple, right axis). F) Purification of RNase1-treated RNA by SPE and re-exposure to RNase 1, with or without previous heat denaturation. Δ: heat.

Given that heat irreversibly affects nicked tRNAs, we studied the effects of phenol-based RNA extraction methods. Because this chemical agent, like heat, can disrupt base-pairing interactions. A recent study of different RNA purification methods for liquid biopsies found that the miRNeasy kit (Qiagen) recovered a broad spectrum of exRNAs associated with different carrier subclasses (Srinivasan et al., 2019). However, when we purified RNase1-treated RNA with this kit following the manufacturer’s instructions, tRNA halves were no longer amenable to enzymatic repair (**Figure S6D**). Similar results were obtained when comparing phenol-free SPE-based purification vs. TRIzol (**Figure 4B**). Interestingly, adding a second SPE-based purification round recovered reparable tRNA halves with a yield close to 100%. Thus, guanidine salts included in the SPE binding buffer do not affect nicked tRNAs, while heat and phenol induce their irreversible separation. As previously observed, the length of the repaired tRNA was slightly shorter (4 – 5 nt) than that of the parental full-length tRNA (**Figure S6E**).

To confirm the effects of phenol and heat by an orthogonal method, we studied RNase-1 treated RNAs by native northern blot (run on TB-Mg^2+^ gels) (**Figure 4C**). Under native conditions, the electrophoretic migration of RNase1-treated tRNA^Gly^_GCC_ was almost identical to that of untreated full-length tRNAs. Only < 10% of the total signal in the treated lane corresponded to *bona-fide* tDRs. This confirms that the rapid disappearance of the full-length tRNA band observed in **Figure 2A** is in fact an artefact caused by denaturing conditions, with > 90% of tRNAs remaining as nicked tRNAs after 1 hour of enzymatic digestion. Interestingly, the migration of nicked tRNAs in native gels was slightly faster than that of full-length tRNAs, consistent with the irreversible loss of the NCCA 3’ overhang inferred from previous assays (**Figure S6E**). Additionally, heat or standard RNA purification methods such as TRIzol or miRNeasy, unlike phenol-free RNA cleanup columns (SPE), induced the irreversible dissociation of nicked tRNAs (**Figure 4C**).

### The 3’ – 5’ RNA ligase RtcB can also ligate nicked tRNAs

RtcB is a ligase involved in tRNA splicing and RNA repair in all domains of life (Englert et al., 2011; Popow et al., 2011; Tanaka and Shuman, 2011). Unlike the 5’ – 3’ T4 Rnl1, RtcB seals broken RNAs with 2′,3′ cyclic phosphate (2,3-cP) and 5′-OH ends (Chakravarty et al., 2012). Thus, RtcB should be sufficient to repair RNase1-treated RNAs without prior end-healing by T4 PNK. Indeed, RtcB regenerated a tRNA-sized band when incubated with RNase1-treated RNAs, and generation of this ligation product was inhibited by heat (**Figure 4D**). Once again, the repaired tRNA-sized band was 4 – 5 nt shorter than the parental full-length tRNA (**Figure S6F**).

### Nicked tRNAs are reverse-transcribed inefficiently unless repaired

Broken phosphodiester bonds could act as roadblocks inhibiting reverse transcription (RT) of nicked tRNAs in their native state, preventing the analysis of nicked tRNAs by RT-PCR or sequencing. To evaluate this possibility, we used a thermostable retroviral reverse transcriptase primed by a gene-specific RT primer aligning to the 3’ end of tRNA^Gly^_GCC_ (**Figure 4E**). Forward and reverse PCR primers were placed at the 5’ and 3’ ends of tRNA^Gly^_GCC_, respectively. Surprisingly, RNase-1 treated RNA was not amplified unless enzymatic repair (T4 PNK + T4 Rnl1) was performed before RT. Consistent with northern blot results (**Figure 3D** and **4A-D**), heating the samples before enzymatic repair also inhibited RT-PCR amplification (**Figure 4E**). Conversely, 5’ tRNA^Gly^_GCC_ halves of 30 nt increased 200 to 700-fold in RNase1-treated samples vs input, illustrating the limitations of small RNA expression analysis in samples containing nicked forms of their parental RNAs.

### Nicked tRNAs are resistant to RNase 1 cleavage in vitro

Having observed that nicked tRNAs can be purified efficiently by SPE, we generated and purified nicked tRNAs and treated them with r-RNase 1 (**Figure 4F**). Nicked tRNAs were not degraded by this treatment (up to 30 min at 37ºC). However, heating and then cooling the nicked tRNAs before exposure to r-RNase1 induced complete degradation of the tRNA halves, now in their single-stranded form. This effect was not induced by heat itself. In conclusion, nicked tRNAs are stable reservoirs of 5’ tRNA halves, which are degradation-prone once dissociated from their 3’ counterparts.

### Nicked tRNAs as a source of intracellular stress-induced tRNA halves

Our enzymatic repair assays were not useful to interrogate the existence of nicked tRNAs inside cells due to the large excess of full-length tRNAs in intracellular samples. We thus resorted to a different strategy based on intracellular RNA fractionation under nondenaturing conditions by size-exclusion chromatography (SEC) (**Figure 5A** to **5D**).

**Figure 5:**
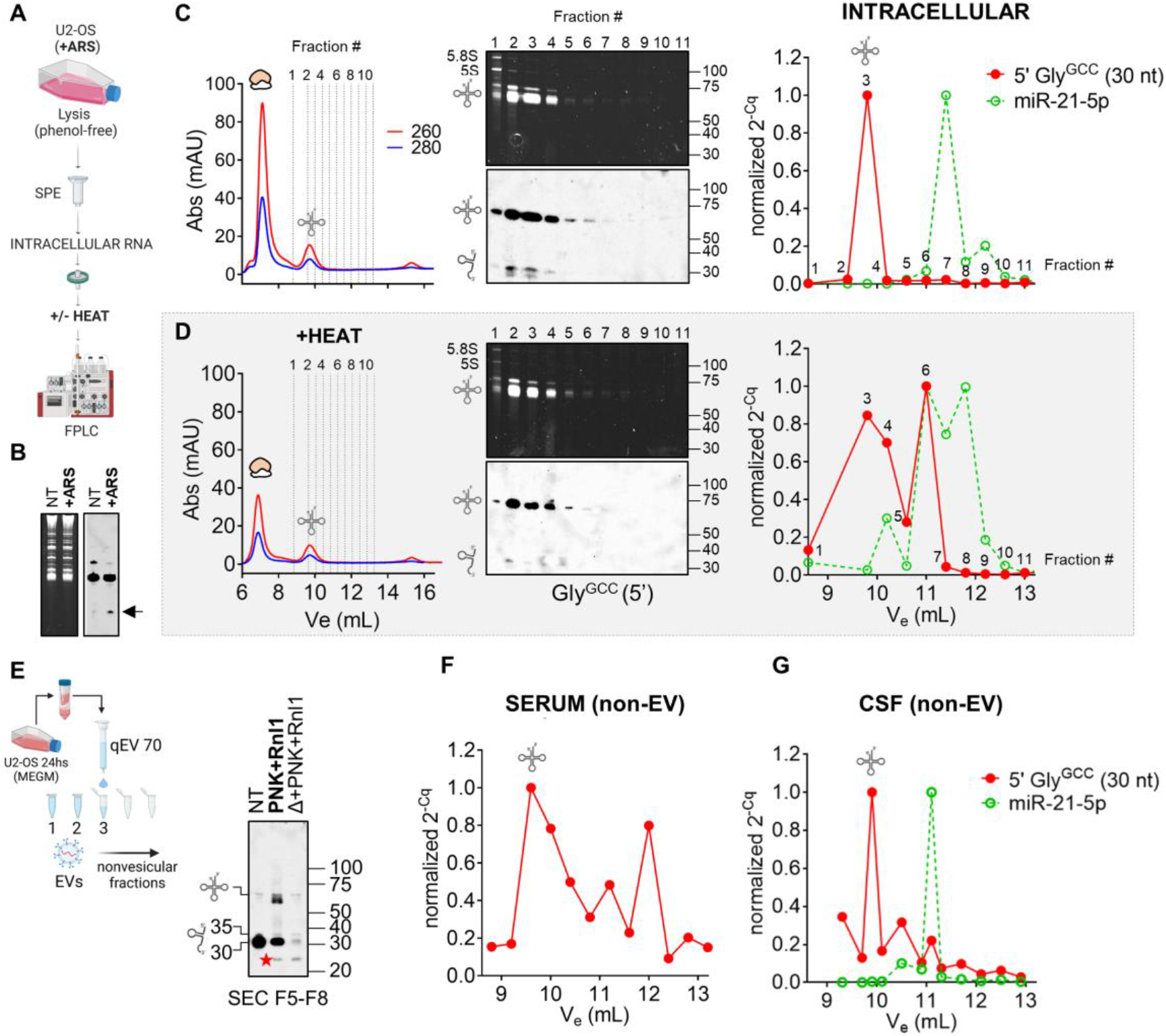
Detection and analysis of nicked tRNAs in intracellular (A-D) and extracellular (E-G) samples. A) Schematic representation of SEC-based (native) intracellular RNA fractionation as performed in (C) and (D). B) Northern blot of intracellular RNAs showing the presence of 5’ tRNA halves in arsenite (ARS)-treated samples. C) Purified RNA from ARS-treated U2-OS cells was separated on a Superdex 75 column using an FPLC system. Selected fractions were analyzed by northern blot (center panel) or by stem-loop RT-qPCR (right panel). C_q_ values were normalized to the fraction containing the highest signal. D) Same as in (C), but the RNA was heat-denatured before injection. E) Cell-conditioned, RI-treated, serum-free medium from U2-OS cells was fractionated with an Izon 70nm qEVoriginal column to prepare EV-depleted fractions. Northern blot analysis of tRNAs & tDRs in pooled nonvesicular extracellular fractions (F5 to F8) before and after enzymatic repair with T4 PNK and T4 Rnl1. Red star: a ∼ 24 nt band that was only observed in the presence of Rnl1. F-G) Separation by SEC (as done in C) of purified RNA from Proteinase K-treated ultracentrifugation supernatants of human serum (E) or CSF (F). Selected eluted fractions were analyzed by SL-RT-qPCR using primers specific for 5’ tRNA^Gly^_GCC_ halves of 30 nt (red) and miR-21-5p (green). A tRNA icon in this figure indicates fractions where full-length tRNAs are expected to elute (if present).

We first treated U2-OS cells with sodium arsenite (Yamasaki et al., 2009) and verified the presence of stress-induced tRNA halves (tiRNAs) by northern blot (**Figure 5B**). Stressed cells were then lysed by phenol-free methods, and intracellular RNA was purified by SPE and separated by SEC using an FPLC system. A tRNA peak was consistently eluted at V_e_ = 9.80 mL, evidenced by the registered absorbance at 260 nm (**Figure 5C**, left panel) and northern blot (**Figure 5C**, center panel). Surprisingly, northern blot bands consistent with 5’ tRNA halves (of 35 and 30 nt) were observed only in the fractions corresponding to the tRNA peak. A more sensitive stem-loop RT-qPCR assay was used to identify tRNA halves in the other fractions (**Figure 5C**, right panel). This assay is specific for mature miRNAs or tRNA halves and does not amplify their precursors (pre-miRNAs or full-length tRNAs, respectively) (Chen et al., 2005; Tosar et al., 2018). While miR-21-5p eluted at V_e_ = 11.4 mL, 5’ tRNA^Gly^_GCC_halves of 30 nt were almost undetectable except at V_e_ = 9.80 mL, where most full-length tRNAs elute. This is consistent with our previous results in non-stressed MCF-7 cells (Tosar et al., 2018). Similar results were obtained for fragments of 35 nt (**Figure S7A**). Heating the RNA before injection (**Figure 5D**) decreased the northern blot signal in the full-length tRNA peak while shifting the main SL-RT-qPCR peak of 5’ tRNA^Gly^ halves to higher elution volumes. Most tRNA-derived fragments of 30 nt co-eluted with miR-21-5p in heated samples.

Overall, while we do not discard that *bona fide* intracellular tiRNAs could be degraded upon cell lysis, our results strongly suggest that at least some (if not most) intracellular tiRNAs are predominantly present in the form of nicked tRNAs.

### Detection of nonvesicular nicked tRNAs in cell-conditioned media

To explore the presence of nicked tRNAs in extracellular samples, we collected cell-conditioned media (CCM, t = 24 hs) from U2-OS cells grown under-serum free conditions and separated EVs from nonvesicular proteins and RNAs by SEC, using commercial qEV columns (**Figure 5E**). To preserve both nicked tRNAs and single-stranded tDRs, we added RNase inhibitors (RI) to the media. As expected in RI-treated serum-free CCM (Tosar et al., 2020), we observed full-length tRNAs in nonvesicular fractions by northern blot (**Figure 5E**; NT lane), albeit most of the signal corresponded to ∼30 nt fragments. In this case, sequential treatment with PNK and Rnl1 generated at least three bands with sizes close to, but slightly shorter than, the size of the parental full-length tRNA. Once again, heating and cooling the samples before ligation abrogated generation of these products, arguing against non-specific ligation in *trans*. The lowest and most intense band had a size of 61 nt, consistent with the main ligation product observed when incubating RNAs in CSF for long periods of time (**Figure S6B**). As previously discussed, this suggests the presence of trimmed forms of nicked tRNAs, lacking both the 3’ NCCA tail and the entire anticodon loop. Once formed, these ligation products could be heated and remained resistant to the processive 3’ – 5’ exonuclease RNase R (**Figure S7B**), suggesting the ligated products are highly structured. In contrast, the remaining 30 – 35 nt band after enzymatic repair was completely degraded by RNase R.

Interestingly, a band of approximately 24 nt was generated by the combined action of PNK and Rnl1 in both heated and nonheated samples (**Figure 5E**; red stars). Because this band was resistant to RNase R (**Figure S7B**) we speculate that it corresponds to a circularized RNA generated by head-to-tail ligation of 5’ tRNA halves. Formation of this product in nonheated samples implies that at least some extracellular tRNA halves are present in fragment form in RI-treated CCM. This suggests that nicked tRNAs are spontaneously separating and releasing single-stranded tRNA halves into the extracellular space. Their detection is facilitated by the low RNase activity in the conditions of this assay.

### Nonvesicular nicked tRNAs circulate in human biofluids

Having validated a method capable of separating nicked tRNAs from single-stranded tRNA halves (**Figure 5C** and **5D**), we implemented this methodology to address the question of whether nicked tRNAs circulate in human biofluids. To do so, small volumes of serum or CSF were diluted in PBS, and EVs were pelleted by ultracentrifugation. RNA was then isolated from Proteinase K-treated supernatants and fractionated by SEC. Surprisingly, 30 nt 5’ tRNA^Gly^_GCC_ halves could be amplified by SL-RT-qPCR (**Figure 5F** and **5G**), mostly in the size range corresponding to full-length tRNAs, which are expected to be absent in these samples (**Figure 2B**).

### Spontaneous uptake of nonvesicular tRNA halves by human epithelial cells

Unless stabilized by dimerization or RNP complex formation, we would expect single-stranded tRNA halves to be degraded in a matter of seconds in extracellular samples with high RNase activity (**Figure 4F**). However, if stable nicked tRNAs diffuse from the sites in which they are generated and they can reach farther cellular populations, their spontaneous dissociation could release single-stranded tRNA halves close to the surface of potentially recipient cells. Could cells then internalize these single-stranded tDRs?

We incubated synthetic and biotinylated 5’ tRNA^Gly^_GCC_ halves of 30 and 35 nt, or scrambled (SCR) versions of these oligonucleotides, with MCF-7 cells grown in serum-free media (**Figure 6A**). No lipids or transfection reagents were used. After 30 min, cells were thoroughly washed, fixed, stained with APC-coupled streptavidin and observed under a confocal microscope. Surprisingly, the red signal of the fluorophore’s emission was observed within cellular boundaries in all conditions where labeled RNAs were included (**Figure 6B and 6C**). As an orthogonal approach, we incubated cells with non-biotinylated SCR oligonucleotides (which do not exist in recipient cells), washed cells thoroughly, and observed a drop of almost 15 PCR cycles when comparing with nontreated cells, based on a SCR-specific SL-RT-qPCR assay (**Figure 6D**). Taken together, these results show that human epithelial cells can spontaneously incorporate single-stranded small RNAs present in the media. Although this uptake route seems to be sequence-independent, most naked extracellular RNAs will degrade before having the opportunity to interact with recipient cells. However, tRNA halves are extremely stable in extracellular samples (**Figure 2**), presumably due to their transport in the form of nicked tRNAs. For this reason, and because their extracellular concentration is increased in the presence of cell stress (**Figure 1D**), we foresee the involvement of extracellular tRNA halves in intercellular communication pathways mediated by naked, nonvesicular RNAs (**Figure 6E**).

**Figure 6:**
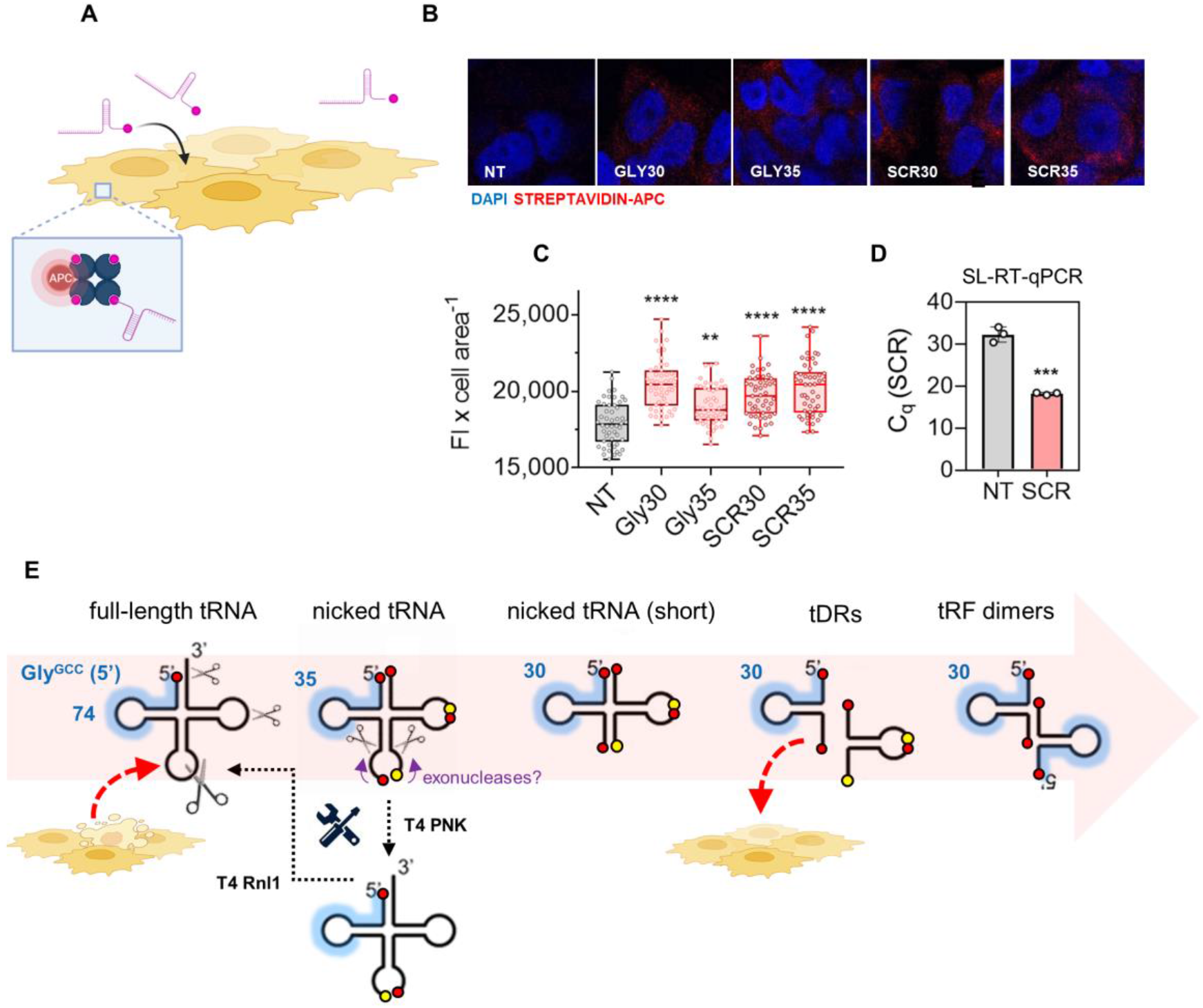
Nicked tRNAs are a source of nonvesicular single-stranded tRNA halves that can be taken up by recipient cells. A) Synthetic 5’ biotinylated RNAs were added to the media of MCF-7 cells and oligonucleotide uptake was confirmed by confocal microscopy using APC-coupled streptavidin (B, C) or by SL-RT-qPCR (D). F) Proposed model.

## DISCUSSION

This work challenges the widespread belief that all RNAs are intrinsically unstable and cannot circulate in extracellular samples unless in the context of RNPs, lipoproteins and/or EVs. Although this holds true for rRNA-derived fragments (**Figure 1**), full-length or nicked tRNAs showed surprisingly long half-lives in human biofluids, even when incubated in their naked forms. While these RNAs might also be present in RNP complexes in the extracellular space, our results demonstrate that protein complexation is not a prerequisite for their remarkable extracellular stability.

An aspect of this study that certainly deserves more attention is the differences in extracellular stabilities observed among tRNAs. We are tempted to speculate that post-transcriptionally modified bases at the anticodon are at least partially responsible for these different behaviors.However, the difference in stability between tRNAs is less pronounced when considering that nicked tRNAs, and not tDRs, are the stable degradation intermediates dictating the abundance of nonvesicular glycine tRNA halves.

Is the full-length tRNA^Gly^_GCC_ unstable in human biofluids? It is efficiently cleaved at several positions by extracellular RNases. So the immediate answer would be “yes”. However, the result of these cleavage events is a molecule that probably still resembles a tRNA, although it bears some broken phosphodiester bonds. Thus, although tRNA^Lys^_UUU_ and tRNA^Gly^_GCC_ show completely different behaviors when analyzed by northern blot after being exposed to RNases, this difference is exaggerated because standard northern blotting forces nicked tRNAs to denature. This is also the case when purifying RNA using phenol and when RNA is heated at any stage in a protocol (e.g., as done during small RNA-seq library preparation). The important conclusion is that we do not always apprehend the true, native form of RNA because most available analytical techniques contain denaturation steps at some point. Additionally, nicked tRNAs cannot be reverse-transcribed and amplified unless repaired or denatured into fragments.

The case of tRNA^Lys^_UUU_ is also interesting. Although it is highly resistant to degradation, once cleaved by r-RNase 1 or by FBS-derived RNase A, it typically does not survive as a nicked tRNA (**Figure 2A**). This implicates extracellular RNases in the degradation of tDRs (Li et al., 2022). In contrast, tRNA^Gly^_GCC_ is sensitive to initial cleavage events, but nicked tRNAs produced therefrom are intrinsically stable. Differential stabilities among distinct tRNA sequences could be considered a new example of non-canonical “moonlighting” functions of tRNAs, what has been related to the diversity of the tRNA isodecoder pool (Avcilar-Kucukgoze and Kashina, 2020).

In bacteria, phage infection (Kaufmann, 2000) or DNA-damage (Hughes et al., 2020) induces regulated tRNA cleavage at the anticodon loop. This process can be reversed by host-encoded tRNA ligases such as RtcB (Tanaka and Shuman, 2011) or by phage antitoxins like T4 PNK and T4 Rnl1. Thus, the existence of nicked tRNAs that can be enzymatically repaired back into full-length tRNAs is well stablished in prokaryotes.

In plants, the phloem sap (PS) has been shown to contain 70-80 nt RNAs, which resemble tRNAs, that are capable of inhibiting translation. Importantly, this activity was lost when purifying PS RNA by TRIzol (Zhang et al., 2009). Thus, plants also seem to generate nicked tRNAs in their extracellular vasculature. The theoretical capacity of nicked tRNAs to act as regulators of the translation machinery was recognized early (Thompson and Parker, 2009). However, nicked tRNAs were not considered as stable degradation intermediates in the subsequent literature (Chen et al., 2021).

Stress-induced tRNA cleavage at the anticodon loop is thought to be an irreversible process. Nevertheless, it should be considered that most of the enzymatic activities used in this work to repair anticodon-cleaved nicked tRNAs are present in human cells (Kroupova et al., 2021; Pinto et al., 2020). Reversibility of stress-induced nicked tRNA formation *in cellulo* arises as an exciting possibility.

In the clinical setting, double-stranded RNAs usually require encapsulation in lipid nanoparticles or GalNAc conjugation for efficient uptake, but single-stranded oligonucleotides are spontaneously endocytosed (Levin, 2019). Thus, our proposed model where nonvesicular nicked tRNAs are carriers of tRNA halves that can transfer information to recipient cells is at least feasible (**Figure 6E**).

On the applied side, we took advantage of phage and bacterial enzymes involved in damaged tRNA repair to develop a new analytical technique that can be used to analyze stable nonvesicular RNAs circulating in biofluids. Although we focused mostly on tRNAs, close inspection of gels shows additional bands that could be restored by combined treatment with PNK and Rnl1 and that were lost in heated samples (**Figure 4B**, yellow stars). This strongly suggests that the nonvesicular RNAome is much more complex than previously thought, in accordance with recent findings (Tosar et al., 2020). Many stable nonvesicular RNAs are single-stranded molecules tightly bound and protected by extracellular RNA-binding proteins (Arroyo et al., 2011; Tosar et al., 2022; Turchinovich et al., 2011). However, other resilient RNAs circulating in biofluids could belong to a “new” category of intrinsically-stable extracellular RNAs (Tosar, 2021). These molecules are predicted to be both highly structured and nicked. There are sequencing methods that can efficiently handle highly structured RNAs (Behrens et al., 2021; Qin et al., 2016). However, nicked RNAs are more challenging because they contain roadblocks for reverse transcriptases (i.e., the broken phosphodiester bonds) and because nicked RNAs are dissociated by phenol or heat. Our enzymatic repair protocol therefore increases the number and types of RNA molecules that can be analyzed and used as disease biomarkers.

## MATERIALS AND METHODS

Detailed versions of these protocols and additional methods are included in ***Supplementary Materials and Methods***.

### Biofluids and cell culture

Biofluids samples were obtained at UC San Diego (CSF) or at the Pasteur Institute of Montevideo (serum, urine), and spun down at 2,000 x g before storage at ≤ -20ºC. Fetal bovine serum (FBS) was from Gibco. Cell-conditioned media (CCM) was obtained from either U2-OS cells grown under serum-free (MEGM, Lonza) or serum-containing conditions (10% FBS), with or without addition of murine RNase inhibitor (RI, NEB). Intracellular tRNA-derived fragments were generated by exposing cells to 500 µM sodium arsenite (Sigma) for 2 hours.

### RNA purification by solid phase extraction (SPE)

RNA was purified by silica-based SPE using Monarch RNA Cleanup Kits (for in vitro assays) or Total RNA Miniprep Kit for intracellular samples (NEB).

### RNA decay assays

For determination of RNA half-lives in biofluids, 1 μg of heated and refolded U2-OS total RNA was incubated at 37ºC for variable periods with 50 μL of undiluted or PBS-diluted biofluid samples. Digested RNAs were purified by SPE and analyzed by northern blot.

### In vitro RNA digestion

For in vitro generation of nicked tRNAs and/or tDRs, 1 μg heated and refolded U2-OS total RNA was incubated with recombinant human RNase 1 (Bon Opus Bio; 0.0625 μg/mL) for 15, 30 or 60 min at 37°C, and purified by SPE.

### RNA ligation assays

In vitro-digested or CCM-derived nonvesicular RNA was incubated for 1 h at 37°C in a 10 μL reaction containing 20 U of RI, 1 mM ATP, 1X T4 PNK reaction buffer, 10 U T4 RNA ligase 1 (Rnl1, NEB) and/or 10 U of T4 PNK (wild-type or 3’ phosphatase minus, NEB). For RtcB ligation, reaction mixtures contained 5 μL of in vitro-digested digested RNA, 20 U of RI, 1X RtcB ligase buffer, 1 mM GTP, 1mM Mn^2+^ and 1 μM RtcB ligase from *E. coli* (NEB).

### Preparation of nonvesicular RNAs from cell-conditioned media

Nonvesicular RNAs from U2-OS cell-conditioned media were separated from EVs by either iodixanol 12-36% density gradients (Jeppesen et al., 2019; Tosar et al., 2020) or by size-exclusion chromatography using 70 nm qEVoriginal columns (IZON).

### Chromatographic fractionation of nonvesicular RNAs from biofluids

EVs were depleted from human serum or CSF samples by ultracentrifugation (256,000 x g and 4°C for 1 h). The supernatants were concentrated by ultrafiltration (10.000 MWCO), treated with phenol-free RNA Binding Buffer (included in RNA Cleanup Kits from NEB) and Proteinase K, and the RNA was purified by SPE. Purified RNAs were then injected in a Superdex 75 10/300 column (GE), using an Äkta Pure FPLC system. Fractions of 0.2 mL were collected and analyzed by stem-loop RT-qPCR.

### Northern blot, RT-qPCR for full-length tRNAs and SL-RT-qPCR for tDRs

DIG-based northern blot and stem-loop RT-qPCR were done as described in (Tosar et al., 2020) and (Tosar et al., 2018), respectively. RT-PCR for full-length tRNAs was done using Superscript IV at 65ºC. Probes, primers, and assay conditions are provided in ***Supplementary Materials and Methods***.

## Acknowledgments

We thank Sergio Bianchi and other members of the Functional Genomics Laboratory (IPMon) and the Analytical Biochemistry Unit (UdelaR). This work was supported in part by the National Institutes of Health (R01GM126150), the National Cancer Institute (NCI) and NIH Office of the Director (UG3/UH3CA241694) and Universidad de la República, Uruguay (CSIC I+D_2020_433). BC received a fellowship from the National Agency of Research and Innovation (ANII, Uruguay; POS_NAC_M_2020_1_163868). J.P.T and A.C. are members of the National System of Researchers (ANII, Uruguay) and the Program for the Development of Basic Science (PEDECIBA, Uruguay).

