## Supplementary material for "Nicked tRNAs are stable reservoirs of tRNA halves in cells and biofluids": Figure S1

### SUPPLEMENTARY FIGURES S1 – S7

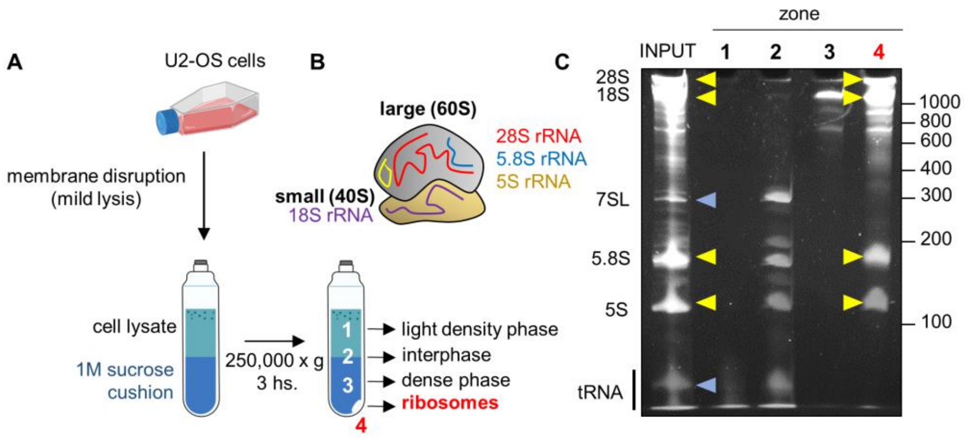


**Supplementary Figure 1:** A) Purification of intracellular ribosomes. B) Schematic representation of an eukaryotic 80S ribosome with the different rRNA species associated with each ribosomal subunit. C) 7M Urea PAGE analysis of purified ribosomes (lane 4) vs. RNAs from the different fractions collected by ultracentrifugation (A). rRNA bands are shown in yellow.

**A**

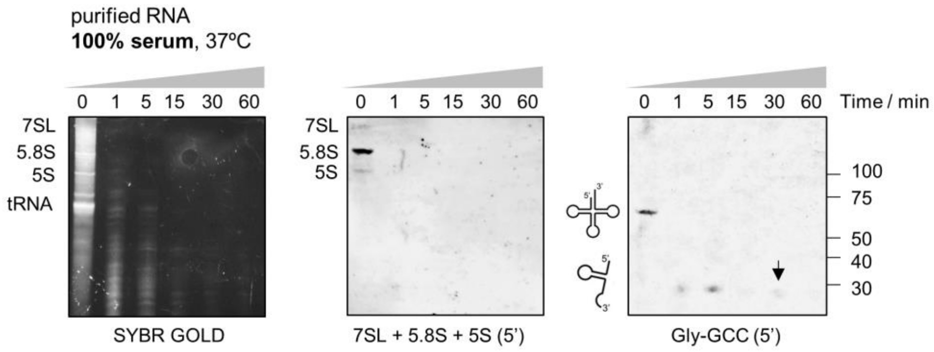


**B**

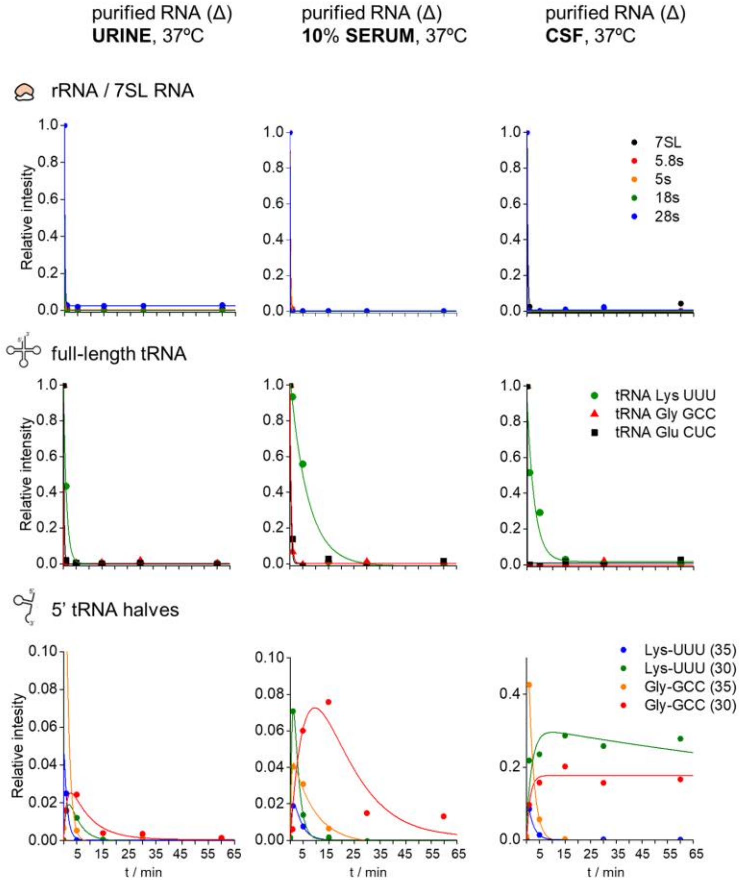


**C**

| BIOFLUID | rRNA 5.8s | 5s | 18s | 28s | tRNA LysUUU | GlyGCC | GluCUC | 5' tRNA halves Lys (35) | Lys (30) | Gly (35) | Gly (30) |
|---|---|---|---|---|---|---|---|---|---|---|---|
| urine | < 6 | < 6 | < 6 | < 6 | 49.5 | < 6 | < 6 | < 6 | 160 | 53 | 327 |
| 10% serum | < 6 | < 6 | < 6 | < 6 | 288 | 15 | 21 | 31 | 75 | 359 | 439 |
| CSF | < 6 | < 6 | < 6 | < 6 | 121 | < 6 | < 6 | 91 | 8866 | 85 | ∞ |

**Supplementary Figure 2:** A) RNA decay in undiluted human serum. T = 0 corresponds to RNAs added to serum and immediately placed on ice. B) Kinetic modeling of RNA decay in human biofluids was done according to the procedure described in supplementary methods. Single exponential decay or transient intermediate models were fitted to band intensities obtained from Figure 2 to determine half-lives for RNA species. C) Half-lives (in seconds) were obtained from kinetic modeling for each RNA in different biofluids.

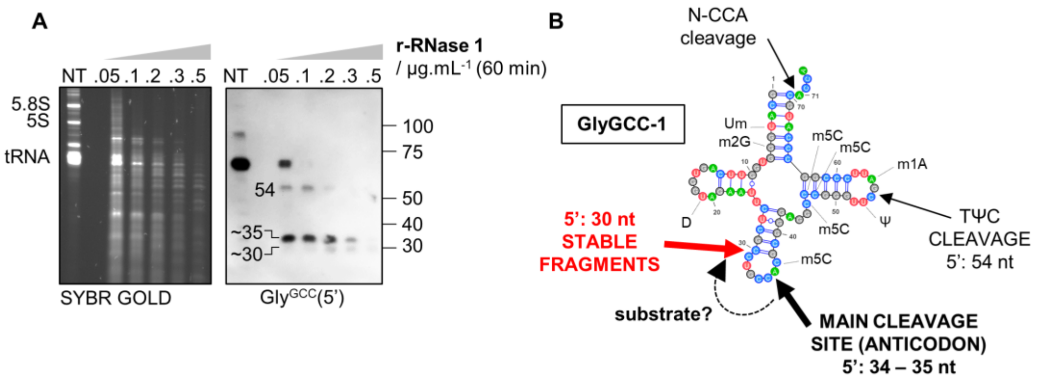


**Supplementary Figure 3:** A) Northern blot using a probe complementary to the 5' end of $tRNA^{Gly}_{GCC}$ after exposure of TRIzol-purified total U2-OS RNA (NT) to different concentrations of recombinant human RNase 1 at 37ºC for 60 min. B) Cloverleaf diagram of human $tRNA^{Gly}_{GCC}$-1 showing modified bases as described in the modomics database (http://genesilico.pl/modomics/) and predicted cleavage sites based on data presented in this study. It is not clear from our results whether cleavage sites at position 30 and 34 – 35 are independent, or whether cleavage at position 34 – 35 is a requisite for efficient cleavage at position 30.

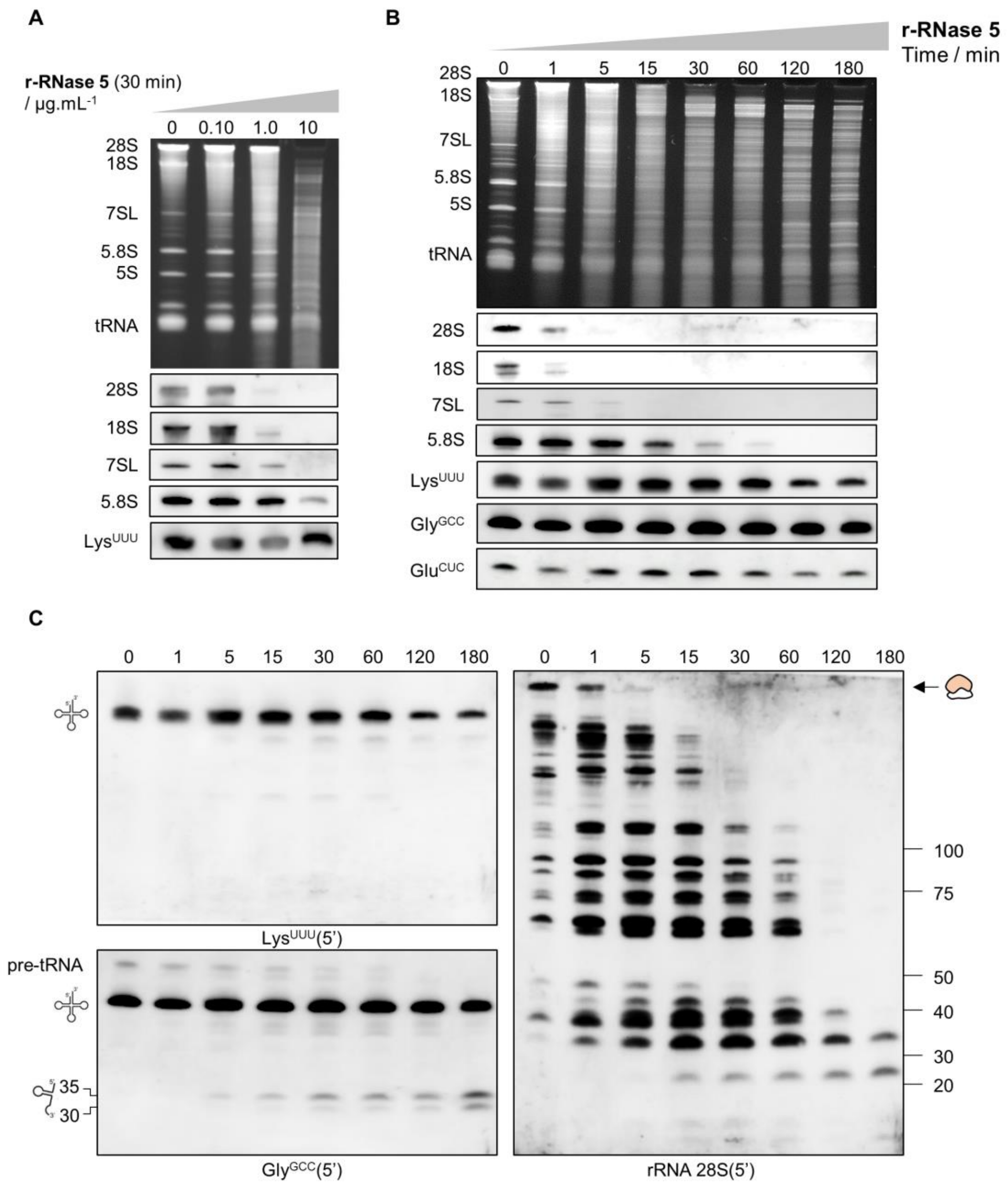


**Supplementary Figure 4:** A) Northern blot analysis of different full-length transcripts after incubating total U2-OS RNA with increasing concentrations of recombinant human RNase 5, for 30 min at 37ºC. B) Same as (A), but the concentration of RNase 5 was fixed and the analysis was performed by varying the incubation time. C) Expanded blots from (B), to show the generation and decay of tRNA-derived fragments (left) and 28S rRNA-derived fragments (right).

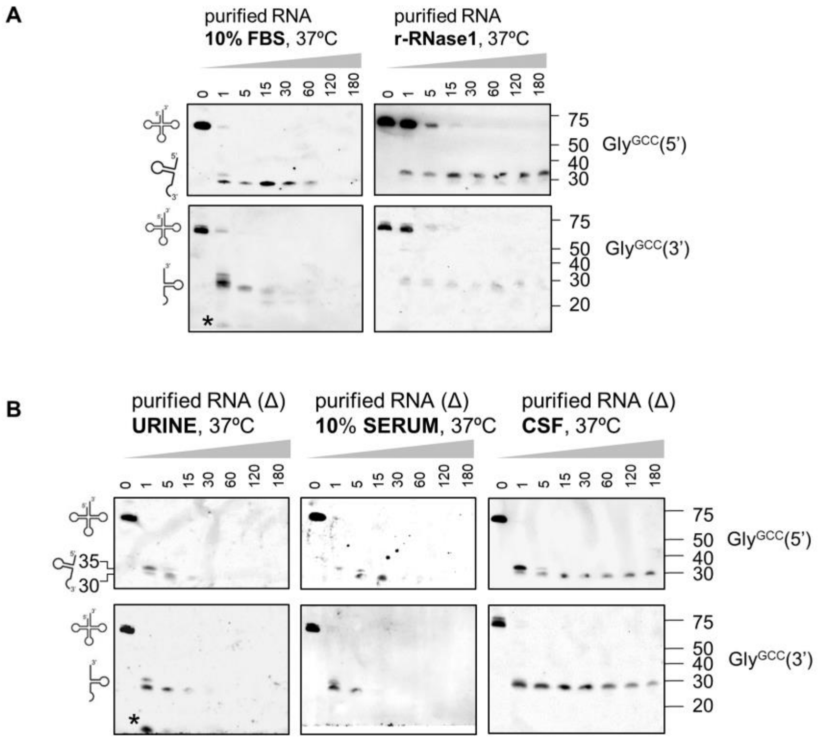


**Supplementary Figure 5:** Northern blots using probes complementary to the 5' and 3' halves of $tRNA^{Gly}_{GCC}$, corresponding to the assays shown in Figure 2, A (A) and Figure 2, B (B).

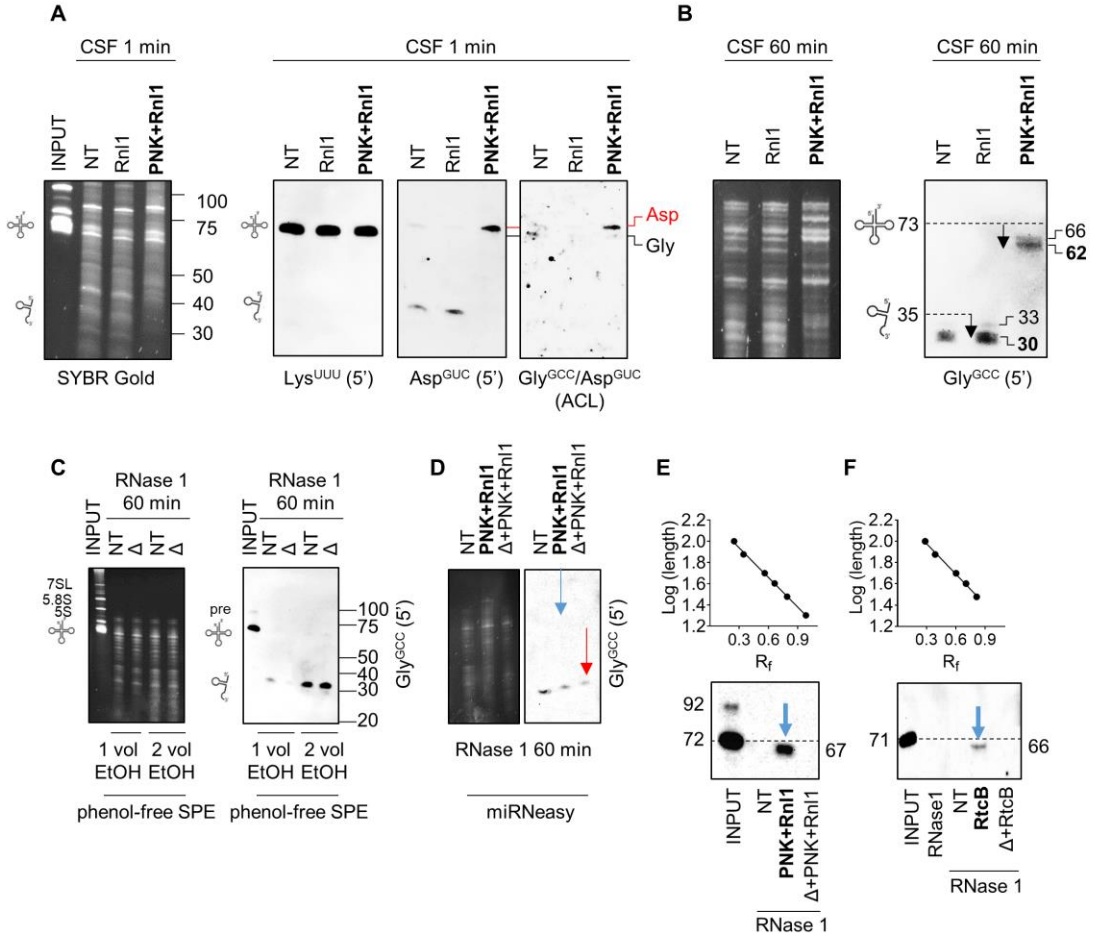


**Supplementary Figure 6:** A-B) incubation of purified RNA in CSF for 1 min (A) or 1 h (B) at 37ºC, followed by enzymatic repair of nicked tRNAs. The northern blot band for tRNA$^{Lys}$$_{UUU}$ serves as a loading control, because tRNA$^{Lys}$$_{UUU}$ is not significantly degraded by short incubations in CSF. The repair of tRNA$^{Asp}$$_{GUC}$ (A, red) is analyzed by using either a 5' (center) or an anticodon-targeting probe (right). NT: fragmented RNA not treated with the enzymatic repair cocktail. C) purification of RNAse1-treated RNA (with or without heat denaturation, Δ) by SPE, following manufacturer's instructions (1 vol EtOH) or duplicating the volume of ethanol added to the binding buffer (2 vol EtOH). D) Enzymatic repair (PNK + Rnl1) of RNase-1 treated RNA purified with the miRNeasy kit. E) Determination of the size of the repaired products in Figure 4, B, based on the $R_f$ method. Migration of the small RNA ladder was used to construct the calibration curve. F) Same as (D), but for Figure 4, D.

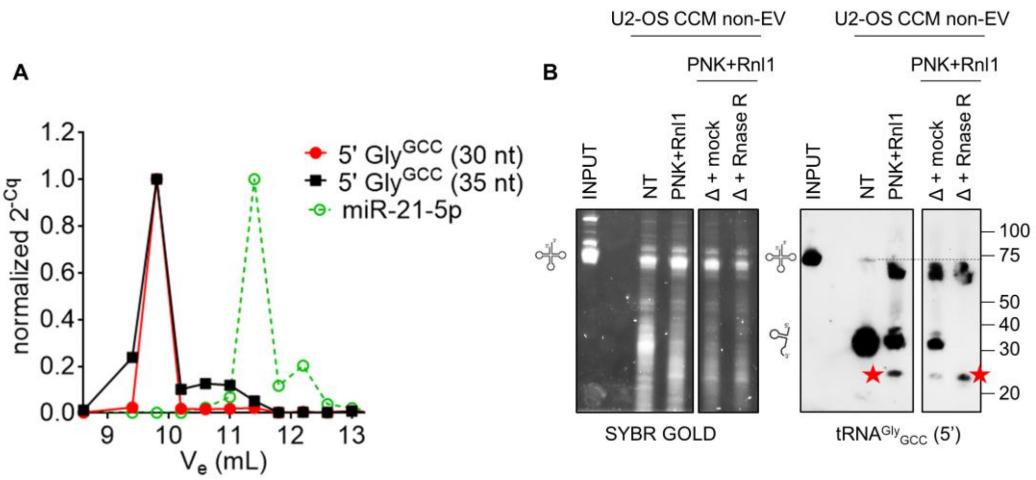


**Supplementary Figure 7:** A) Data from Figure 5, C, but overlapping the signal obtained when using primers specific for tRNA$^{Gly}_{GCC}$ 5' fragments of exactly 35 nt (black). B) A replicate of the assay shown in Figure 6, B, but the ligated RNA was now heated after PNK + Rnl1 treatment (Δ), cooled down to room temperature, and incubated with RNase R to degrade single-stranded linear and unstructured RNAs. Red stars: short, RNase R-resistant ligation products, presumably corresponding to circularized single-stranded 5' tRNA$^{Gly}_{GCC}$ halves.
