## Supplementary Materials and Methods for "Nicked tRNAs are stable reservoirs of tRNA halves in cells and biofluids"

**Biofluid samples**

CSF was obtained from a patient with progressive supranuclear palsy at the Movement Disorder Center of the University Of California San Diego School Of Medicine, following approved procedures and protocols. Samples were centrifugated at 2,000 x g for 10 min at room temperature before freezing at -80ºC.

Blood and urine samples were collected at the Pasteur Institute of Montevideo. Blood samples were obtained by venipuncture from a healthy donor in serum blood collection tubes and centrifuged at 2,500 rpm for 15 min to separate serum, which was stored at -20ºC until use.

20 mL of a human urine sample was retrieved within 1 h of collection from a healthy donor in a sterile container. The sample was transferred to a 10 mL Falcon tube and centrifuged at 300 x g and 4°C for 10 minutes, followed by centrifugation at 2000 x g and 4°C for 20 minutes. The supernatant was then stored at -20°C until use.

**Cell culture**

U2-OS cells were cultured in DMEM (Gibco) with 4.5 g/L D-glucose and 110 mg/L sodium pyruvate, supplemented with 10% fetal bovine serum (FBS) (Gibco), without antibiotics. For assays performed under serum-free conditions, cells were washed and incubated in MEGM (Lonza), without the inclusion of bovine pituitary extracts. When indicated, 2 µL (80 U) recombinant murine RNase inhibitor (RI, New England Biolabs) were added to fresh media.

**Generation of stress-induced tRNA halves and intracellular RNA purification**

For the generation of stress-induced tRNA halves (tiRNAs), 500 µM of freshly made sodium arsenite (Sigma-Aldrich) were added to U2-OS cells grown to 90% confluence in DMEM + 10% FBS. After 2 h at 37°C, the medium was removed and cells were washed with warm 1X PBS prior to RNA extraction using a Total RNA Miniprep Kit (NEB). Manufacturer’s instructions were followed, except for two modifications: i) cell lysis was performed by adding 2 mL of lysis buffer directly to the cell monolayer, followed by a short (5 min) incubation at room temperature; and ii) twice the indicated volume of ethanol was added to the RNA sample after elution from the DNA-binding column.

**Purification of intracellular ribosomes**

Ribosomes were purified from U2-OS cells following published methods with modifications [1,2]. Briefly, U2-OS cells were cultured in T75 flasks with DMEM supplemented with 10% FBS and no antibiotics. When cells reached 95-100% confluency, CCM was discarded and cells were washed three times with DPBS (without $Ca^{2+}$ or $Mg^{2+}$) to remove any trace of RNAse activity. Cells were detached with a scraper, suspended in DPBS, and pelleted by centrifugation (10min, 500g, 4°C). Pelleted cells were gently resuspended in 2ml of mild lysis buffer (20mM Tris-Cl, pH=7.4; 300mM NaCl; 6mM $MgCl_2$; EDTA-free protease inhibitor cocktail (Roche); 200U murine RNase inhibitor; and 0.5% v/v IGEPAL CA-630 (Sigma-Aldrich)) and kept on ice for 25 min to release their cytoplasmic content. Unlysed cells, nuclei, and mitochondria were removed by sequential centrifugation at 750 x g and 12,500 x g for 10 min at 4°C. The remaining supernatant was layered on top of a 1M sucrose cushion prepared in purification buffer (50mM Tris-Cl, pH=7.4; 150mM KCl, 5mM $MgCl_2$, 200U murine RNase Inhibitor) and centrifuged for 3hs at 250.000g and 4°C using a Sorvall Discovery M120 SE with a S100-AT6 rotor. After removing the supernatant, a translucent pellet containing purified ribosomes was clearly visible. The pellet was washed two times with 300 µl of ice-cold ultrapure RNase-free water, resuspended in ice-cold working buffer (50mM Tris-Cl, pH=7.4; 25mM KCl; 5mM $MgCl_2$) and stored at -80°C until use. Ribosome quality was assessed by RNA polyacrylamide gel electrophoresis under denaturing conditions.


1. Khatter, H. *et al.* Purification, characterization and crystallization of the human 80S ribosome. *Nucleic Acids Res* **42**, e49–e49 (2014).

2. Belin, S. *et al.* Purification of Ribosomes from Human Cell Lines. *Curr Protoc Cell Biology* **49**, 3.40.1-3.40.11 (2010).


**RNA purification by solid phase extraction (SPE)**

SPE was carried out using Monarch RNA Cleanup Kits (10 µg binding capacity columns, NEB) except for the purification of stress-induced tRNA-derived fragments from U2-OS cells, where the Total RNA Miniprep Kit (NEB) was used. Purification was carried out following manufacturer's instructions, but in several experiments, the volume of ethanol added to the binding buffer was doubled to avoid small RNA loss.

**RNA decay assays and in vitro RNA digestion**

To study the effect of proteins on rRNA stability, 25 µg total RNA (purified from U2-OS cells by TRIzol extraction) or 3 µg of rRNA present in purified ribosomes from U2-OS cells, were incubated in 600 µL RPMI containing 10% FBS, with or without the addition of 200U of RI. 100 µL aliquots were taken at predetermined times, and total RNAs contained in these aliquots were extracted after mixing with 900 µL of TRIzol (ThermoScientific).

For determination of RNA half-lives in biofluids, 1 µg of heated and refolded (1 min at 90°C followed by 30 min at room temp.) U2-OS total RNA was added to 50 µL undiluted human CSF, undiluted human urine, undiluted human serum, diluted human serum or FBS (both 10% in 1X PBS). Additionally, 1 µg of heated and refolded total RNA was incubated at 37ºC for variable periods with recombinant human RNase 5 at 55 µg/mL (r-RNase 5; R&D Systems) or recombinant human RNase 1 at 0.5 µg/mL (r-RNase 1; Bon Opus Bio). Reactions were stopped by the addition of SPE binding buffer and RNA cleanup. In all cases, samples were analyzed by northern blot.

For in vitro generation of nicked tRNAs and/or tDRs, 1 µg heated and refolded U2-OS total RNA was mixed with 40 µL of r-RNase 1 (Bon Opus Bio; diluted in PBS at 0.0625 µg/mL) for 15, 30 or 60 minutes at 37°C. For RNA digestion in human CSF, 1 µg heated and refolded U2-OS total RNA was added to 10 µL of CSF, and incubated for 1 or 60 min at 37°C.

**Kinetic analysis of RNA decay**

The half-lives of the RNA fragments in different environments were obtained by fitting a single exponential function or a model for an intermediate in consecutive first-order reactions to the intensities of the northern blot bands *vs* time [3]. If the decay of the original full-length RNA band could not be observed within the first incubation minute, a half-life of < 6 s was assigned.

A single exponential model (Intensity = Amp*exp(-k*t) + Y, where Amp is the amplitude, t is time, k is the first-order rate constant and Y is an offset value) was fitted to the decay of the initial full-length RNA (when possible) or to the first degradation product (when the decay of the parent full-length RNA was immediate). Transient intermediates that build up and later fall were considered to follow an irreversible model A → B → C. A function (Intensity = (k1*Amp)/(k2-k1)*(exp(-k1*t)-exp(-k2*t)) + Y, where k1 is the first-order rate constant for the decay of the parent fragment A and k2 is the constant for the decay of fragment B, was fitted to the band intensities to obtain k2. Half-lives were calculated as $t_{1/2} = \ln(2)/k$. OriginPro 8.6 was used to analyze the data.

3. Espenson, J. Chemical Kinetics and Reaction Mechanisms 2nd Edition, (1994)

**Northern Blotting**

For denaturing northern blots, 5 µL of RNA were mixed with 5 µL of loading buffer containing 95% formamide, 1mM EDTA, 0.02% SDS, 0.02% bromophenol blue and 0.01% xylene cyanol, heated to 65ºC for 5 min and ran in 10 x 10 cm 10% polyacrylamide gels containing 7M urea in 1X Tris-borate EDTA (TBE, pH 8.4). For non-denaturing northern blots, the RNA samples were mixed with 1 µL of 6X native loading buffer and run in gels containing 1X Tris-borate (pH 8.3) and 10 mM $MgCl_2$ (TB + $Mg^{2+}$) gels. Gels were run for 80 minutes in 0.5X TBE or 0.5X TB + $Mg^{2+}$ running buffer at room temperature, stained with 1X SYBR gold (Invitrogen) and then transferred to positively charged nylon membranes (Roche) using a semi-dry Trans-Blot Turbo Transfer System (Bio-Rad) in 0.5X TBE at constant I = 0.3 A for 30 minutes.

Membranes were UV cross-linked and hybridized for 16 h at 42°C with digoxigenin-labeled DNA probes in DIG Easy Hyb solution (Roche). After hybridation, membranes were washed for 5 minutes at room temperature with low stringency wash buffer (twice, 2X SSC/0.1% SDS), 5 minutes at 42°C with high stringency wash buffer (1X SSC/0.1% SDS), blocked for 30 minutes at room temperature with 1X blocking solution (Roche) and probed for 30 minutes with an alkaline phosphatase-labeled anti-digoxigenin antibody (Roche). Membranes were washed twice with 1X TBS-T for 5 min and then incubated in detection buffer (Roche). Signals were then visualized with CDP-Star, ready-to-use (Roche) and detected using an Amersham ImageQuant 800 imager (GE Healthcare/Cytiva).

Probes (5' to 3'):

rRNA 28S 5': CACGTCTGATCTGAGGTCGC
rRNA 18S 5': ATGCTACTGGCAGGATCAAC
rRNA 5.8S: CGCACGAGCCGAGTGATCCAC
rRNA 5S 5': GGTGGTATGGCCGTAGAC
7SL: CACTACAGCCCAGAACTCCTGGACT
$tRNA^{Gly}_{GCC}$ 5': CTACCACTGAACCACCCATGC
$tRNA^{Lys}_{UUU}$ 5': CTGATGCTCTACCGACTGAGCTATCCGGGC
$tRNA^{Asp}_{GUC}$ 5': TCACCACTATACTAACGAGGA
$tRNA^{Glu}_{GUC}$ 5': TAACCACTAGACCACCAG
$tRNA^{Gly}_{GCC}$ 3': GCCGGGAATCGAACCCGGGCCTCCCGCG
$tRNA^{Gly}_{GCC}/^{Asp}_{GUC}$ ACL: TCCCGCGTGGCAGGCGAGAA
$tRNA^{Lys}_{UUU}$ ACL: CCTCAGATTAAAAGTCTGATG

All oligonucleotides were obtained from Integrated DNA Technologies (IDT, USA), and labeled with DIG Oligonucleotide Tailing Kit, 2nd generation (Roche), following manufactuer's instructions.

**Stem-loop RT-qPCR for tDRs.**

Specific cDNA of $tRNA^{Gly}_{GCC}$ 5' halves and miR-21-5p was obtained with SuperScript II (ThermoScientific) based on the stem-loop RT-qPCR method, as previously described (Tosar et al. 2018). Purified RNAs were heated at 90°C for 1 minute and immediately placed on ice before reverse transcription. qPCR was performed using a QuantStudio 3 Real Time PCR System (ThermoScientific) with FastStart Universal

SYBR Green Master (Rox; Roche). $2^{-Cq}$ values were obtained and normalized against the fraction containing the highest signal.

Primers (5' to 3')

Stem-loop RT primer (“X” denotes assay-specific 3’ overhangs):

GTCGTATCCAGTGCAGGGTCCGAGGTATTCGCACTGGATACGACXXXXXX
$tRNA^{Gly}_{GCC}$ (5' half, 35 nt, 3' overhang): GGCAGG
$tRNA^{Gly}_{GCC}$ (5' half, 30 nt, 3' overhang): GCGAGA
miR-21-5p (3' overhang): GTCAAC

$tRNA^{Gly}_{GCC}$ (5' half, F-primer): CCGCATTGGTTCAGTGGT
miR-21-5p (F-primer): gccccgTAGCTTATCAGACTGATGT
$9^{GG/AA}$ (F-primer): gctcgGCATTGGTAATTCAGTGGTA
Universal reverse primer: GTGCAGGGTCCGAGGT

Lower case letters indicate added bases to increase melting temperature.

All primers were obtained from Integrated DNA Technologies (IDT, USA).

**RT-PCR for full-length tRNAs.**

2 µL of 10 ng/µL U2-OS total RNA (input) or RNase 1-treated RNAs (starting from an equivalent amount of input RNA) were mixed with 1 µL of 2 µM gene-specific RT primer and 1 µL of 10 mM (each) dNTP mix in a reaction volume of 11 µL, incubated for 5 min at 65°C, and chilled on ice for at least 1 min. Annealed RNAs were mixed with 4 µL 5X SuperScript IV (SSIV, Thermo) buffer, 1 µL 0.1M DTT, 2U RI and 20U SSIV RT. The reaction was incubated for 10 min at 65°C and then for 10 min at 80°C. The cDNA was diluted 1/2 prior to qPCR or 1/10 for endpoint PCR.

Gene-specific RT primer for $tRNA^{Gly}_{GCC}$1-4: GCGTCTCACTTATGCACAGCGAACT**TGCATGGGCCGGG**

$tRNA^{Gly}_{GCC}$-1 (F-primer): GCATGGGTGGTTCAGTGGTA
Universal tRNA reverse primer: GTCTCACTTATGCACAGCGAA

**RNA ligation assays**

For ligation assays involving T4 RNA ligases, 5 µL of in vitro-digested RNA (or RNA purified from CCM) were incubated for 1 h at 37°C in a 10 µL reaction containing 20U of RI, 1mM ATP, 1X T4 PNK reaction buffer, and desired enzyme combinations. Enzymatic cocktails included: 10U T4 RNA ligase 1 (Rnl1, NEB) or T4 RNA ligase 2 (Rnl2, NEB) and/or 10U of T4 PNK (wild-type or 3' phosphatase minus, NEB). As a control, RNA was heat-denatured before enzymatic treatment (1 min at 90°C and immediately placed on ice).

For RtcB ligation, reaction mixtures (10 µL; 1 h at 37°C) contained 5 µL of in vitro-digested digested RNA (with or without previous heat denaturation), 20U of RI, 1X RtcB ligase buffer, 1 mM GTP, 1mM $Mn^{2+}$ and 1 µM RtcB ligase from *E. coli* (NEB). The reactions were stopped by addition of 2X RNA Loading Dye and analyzed by northern blot.

**RNase R treatment**

RNA from CCM treated with T4 PNK + T4 Rnl1 was purified by SPE. Reaction mixtures containing 10 µL of ligated RNAs with or without heat denaturation, 1X RNase R Reaction buffer and 20U of RNase R (Lucigen), in a 20 µL final volume, were incubated for 1 h at 37°C. Reactions were stopped by the addition of 100 µL of the RNA Binding buffer included in SPE kits.

**Analysis of nonvesicular RNAs by density gradient separation**

U2-OS cells were incubated with fresh complete medium (DMEM + 10% FBS, both Gibco) in a T150 culture flask until 90% confluency. Fresh medium was added containing or not 500 µM sodium arsenite (Sigma-Aldrich) for 2 hours. Cells were then washed with 1x DPBS (Gibco) and incubated in complete medium for 6 additional hours. The CCM was centrifuged at 300 x g and then at 2,000 x g at 4ºC (20 min) and concentrated to 1 mL by ultrafiltration, using Vivaspin centrifugal filters (Sartorius) with a cut-off of 10 kDa. The CCM was then loaded on the bottom of a 12 – 36 % iodixanol gradient prepared as described in Tosar et al. (2020) and centrifuged overnight at 150,000 x g and 4ºC (SW 40 Ti rotor, Beckman Coulter) using an Optima XPN ultracentrifuge (Beckman Coulter). 12 fractions of 1 mL were collected from the top of the tube. Fractions >7 are EV-depleted (Tosar et al. 2020; Jeppesen et al. 2019). RNA from a volume equivalent to 50% of each fraction was extracted by Trizol LS (ThermoScientific), following manufacturer's instructions, and analyzed by northern blot.

**Fractionation of cell-conditioned media by SEC (IZON columns).**

U2-OS cell-conditioned medium (CCM) obtained in MEGM + 80U RI (1x T75 flask at 80% confluency, conditioning time = 24 h) was harvested, centrifuged at 2,000 x g, and concentrated to 500 µL by ultrafiltration, using Vivaspin centrifugal filters (Sartorius) with a cut-off of 10 kDa. The concentrated CCM was then fractionated using a 70 nm qEVoriginal column (IZON) and an IZON automatic fraction collector, using PBS as the mobile phase. All procedures were done at 4ºC. The columns were pre-chilled by running 1 column volume of chilled PBS before injecting the samples. Thirteen fractions of 1 mL were collected after the void volume had passed through the column. Fractions #1 and #2 contained most EVs and were analyzed in parallel for other projects. EV-depleted fractions #5 to #8 were isopropanol-precipitated, pooled, and the RNA was purified from these fractions by SPE.

**Identification of nonvesicular nicked tRNAs in biofluids**

A 200 µL healthy donor human serum sample was thawed and centrifuged at 2,000 x g and 4°C for 10 min and diluted in 12 mL of PBS. EV depletion was performed by ultracentrifugation at 256,000 x g and 4°C for 1 h in an Optima XPN ultracentrifuge (Beckman Coulter) with a SW 40 Ti rotor. The supernatant was then concentrated by ultrafiltration to 200 µL, using 10.000 MWCO Amicon Ultra-15 Centrifugal filters (Merck). Then, 440 µL of the RNA Binding Buffer included in RNA Cleanup kits (NEB) was added to the concentrated supernatant and treated with 20 µL Proteinase K (QIAgen) for 30 minutes at 37°C. Nucleic acids were then purified by SPE and eluted in 50 µL nuclease free $H_2O$ (Invitrogen). An identical procedure was followed for CSF (input: 400 µL). Samples were analyzed by size-exclusion chromatography using an FPLC system.

**Size exclusion chromatography (FPLC)**

Nonvesicular samples from human serum and CSF, or total RNA from U2-OS cells stressed with sodium arsenite (with or without heat denaturation and refolding) were diluted in 1X PBS (500 µL) and centrifuged at 10.000 x g and 4°C for 10 min prior to injection into a Superdex 75 10/300 column (GE). Size-exclusion chromatography (SEC) was performed at 0.5 mL/min in 0.2 µm-filtered 1X PBS with an Äkta Pure FPLC system and 0.2 mL fractions were collected while monitoring the absorbance at 260 and 280 nm. Nucleic acids from selected fractions were ethanol-precipitated (700 µL absolute anhydrous ethanol; 100 µL 3M NaAc, pH = 5.2; 0.5 µL Glycogen Blue) overnight at -20ºC and centrifuged at 12.000 x g and 4ºC for 15 minutes. The pellet was washed with 500 µL 75% ethanol, centrifuged at 12.000 x g and 4ºC for 15 minutes and resuspended in 10 µL of nuclease-free water (Invitrogen) before northern blot or stem-loop RT-qPCR.
